# Estimation of the time course of excitatory and inhibitory conductance during oscillatory periods

**DOI:** 10.64898/2026.08.10.743856

**Authors:** R.M. Delicado-Moll, A. Guillamon, A.E. Teruel, C. Vich

**Affiliations:** Departament de Matemàtiques i Informàtica and Institute of Applied Computing and Community Code (IAC3), Universitat de les Illes Balears, Palma, Illes Balears, Spain; Departament de Matemàtiques, Universitat Politècnica de Catalunya, Barcelona, Catalunya, Spain

## Abstract

Determining the amount of information a neuron receives per unit of time is key to understanding brain connectivity and how neural networks encode and transmit information. In particular, estimating this information flow by distinguishing between excitatory and inhibitory synaptic contributions is critical to understanding neural network function, as maintaining the excitation-inhibition (E/I) balance regulates neuronal excitability and circuit stability, whereas its disruption can lead to a plethora of brain disorders, including neurodegenerative and psychiatric conditions. However, because synaptic conductances cannot be measured directly, inverse methods are required to infer them from the membrane potential —a readily measurable quantity. Although partial solutions have been proposed, accurately estimating these conductances remains a significant challenge due to the complexity and diversity of the inputs. This is particularly true in the spiking regime, where neurons actively fire.

In this work, we introduce a novel computational strategy that combines two critical metrics extracted from the time course of the membrane potential recording: the amplitude of the spike and the interspike interval. By using these quantities, the proposed method enables the accurate separation of excitatory and inhibitory contributions, yielding highly favorable results in the spiking regime.

**Author summary:** Quantifying the continuous stream of inputs a neuron receives is key to understanding brain connectivity. Inside the brain, individual cells must maintain a tight balance between excitation and inhibition (E/I) to process information correctly, as any disruption in this equilibrium can impair its functionality. However, directly measuring the underlying excitatory and inhibitory synaptic conductances is technically challenging, and existing mathematical tools often fail when neurons enter their active firing regime.

In this work, we introduce a novel computational strategy designed to extract and separate these time-varying conductances directly from the neuron’s spiking activity. By dynamically tracking just two accessible metrics – the amplitude of the spikes and the time intervals between them – our algorithm estimates both conductance profiles with high precision. Furthermore, we demonstrate that this procedure is highly robust against realistic experimental noise and data variability, providing an accessible framework that does not require complex hardware or an unfeasible number of repetitive experimental trials. By tracking changes in the E/I ratio of the synaptic input, this method provides an efficient approach to detecting pathological imbalances and understanding how local connectivity shapes cellular functionality.

## Introduction

Functional connectivity is widely recognized as a crucial feature for understanding brain dynamics. While most research focuses on large-scale connectivity, estimating the local inputs that individual neurons or neuronal populations receive remains at the core of the problem. The activity of presynaptic neurons is channeled through synaptic conductances, which can be either excitatory (*g*_*E*_(*t*)) or inhibitory (*g*_*I*_ (*t*)).

How the continuous stream of synaptic inputs arriving at a single neuron evolves over time remains an open question, relevant for deciphering brain plasticity and cellular functionality. At the single-cell level, individual neurons can receive tens of thousands of inputs, whose ratio must remain tightly balanced over millisecond timescales [1–3]. Disruption of this instant-to-instant excitatory/inhibitory (E/I) conductance balance causes severe neuronal deficits. These cellular-level disruptions underlie various neurodegenerative diseases, such as Alzheimer’s [4] and Parkinson’s disease [5, 6], as well as neurodevelopmental disorders like autism [7]. Furthermore, this dynamic E/I balance is tightly involved in learning ability, during which distinct synaptic plasticity rules operate locally across different dendritic compartments [8]. Consequently, tracking and quantifying these precise, time-varying synaptic changes at high temporal resolution is essential to understanding how individual cells compute and exploit fast E/I shifts for input gating during complex behaviors [9].

A prominent example of this requirement is found in stellate cells, a class of neurons whose precise E/I estimation is critical for cognitive and motor computations. In the medial entorhinal cortex (MEC), layer II stellate cells are fundamental for spatial navigation, acting as grid cells, head-direction cells, or speed cells that generate the brain’s internal GPS [10]. Accurate tracking of the E/I conductance balance in these cells is essential to reveal how they integrate path-integration signals and self-motion cues without losing spatial tuning or causing drift in location estimation [11]. Similarly, in the context of motor coordination, stellate cells in the cerebellar molecular layer exert tight inhibitory control over Purkinje cells [12]. Quantifying the time-varying E/I shifts arriving at these cerebellar interneurons is important to deciphering how the brain fine-tunes real-time motor commands, error correction, and the learning rules that sustain smooth closed-loop control.

The main concern on the estimation of the conductances is the problem to extract them from the activity of the brain in *in vivo* experiments. Direct measurements of synaptic conductances are technically challenging, particularly in vivo and in behaving animals. Consequently, many studies rely on inverse methods to infer excitatory and inhibitory conductances from neural recordings. When attempting to estimate conductances from an observable (typically the membrane potential, *V* (*t*)), one major problem in designing inverse methods is the nonlinearity of the frequency-input current (*f* − *I*) curve due to underlying oscillatory mechanisms that induce spikes in the membrane potential of the receiving neuron. Moreover, when observable data are extracted from the neuron, they are subject, on the one hand, to possible measurement errors, and on the other hand, to data variability if multiple trials of the same experiment are required. Finally, another complication is that there is no a universal neuronal model to describe the activity of different neurons, making it even more difficult to design estimation procedures that are valid for a wide range of neuron types.

Several methods to estimate synaptic conductances have been proposed in the literature. Early approaches assumed that the membrane potential remains constant over short time intervals and that nonlinear ionic currents are negligible. These assumptions linearize the estimation problem, making the estimation of conductances mathematically feasible. Some of these techniques are based on deterministic models [13–17], whereas others employ stochastic frameworks to improve estimates by taking into account possible sources of noise [18–21]. However, [22] demonstrated that assuming such linearity yields highly inaccurate estimates when neuronal activity enters the spiking regime. Consequently, these traditional methods have been restricted to membrane potential traces exhibiting exclusively subthreshold activity, or to recordings where filtering techniques are applied beforehand to eliminate sporadic spikes. Subsequently, it was also shown that linearity can also be lost if nonlinear ionic currents are activated within the subthreshold regime [23], though specific solutions have been developed to overcome this particular limitation [23, 24].

Concerning solutions to cope with nonlinearities in the spiking regime, new estimation procedures to estimate the synaptic conductances have been designed [25–27]. In [25], mathematical insights are provided to estimate the synaptic conductance time course using previous knowledge of the relation between the total synaptic conductance and the interspike period. However, this study only presents a proof of concept in a piecewise linear toy model to predict the total synaptic conductance. Afterwards, [27] extended this idea to adapt the procedure to more realistic neural models, applying the technique to a pyramidal cell model, yet without addressing the challenge of isolating excitatory from inhibitory conductances. Following the same ideas, [26] uses firing-clamp techniques to extract, from different trials, a relation between the excitatory conductance, the inhibitory one, the period of oscillation and the amplitude of the spikes. Although this new technique improves with respect to the previous ones, given the fact that it is capable to discern between excitation and inhibition, it presents the disadvantage that it requires a large number of trials extracted directly from the target neuron, which may present unforeseen errors coming from intertrial variability and noise measurements. Also, this method requires having the appropriate instruments to carry out this type of experiment.

In this paper we develop an estimation procedure based on the techniques presented in [26] and [27]. In particular, using a base model capable to reproduce the membrane potential of a target neuron, from which we want to estimate the excitatory and inhibitory conductance time course, we extend the estimation procedure presented in [27] by considering both the amplitude of the membrane potential peaks and the spike frequency. We computationally extract a relation between constant synaptic conductances (*g*_*E*_, *g*_*I*_), the oscillatory period *T* (the inverse of the spike frequency) and the amplitude *A*; by applying it dynamically to the membrane potential time course, we estimate constant conductances within sufficiently small time intervals, yielding a discretized estimate of their time course that is subsequently refined by interpolation.

In Figure 1, we present a sketch on how the procedure is designed (green pathway), how do we test the estimation method (blue pathway), and finally, how to use the procedure to estimate the conductances of experimental data (pink pathway).

**Fig 1.**
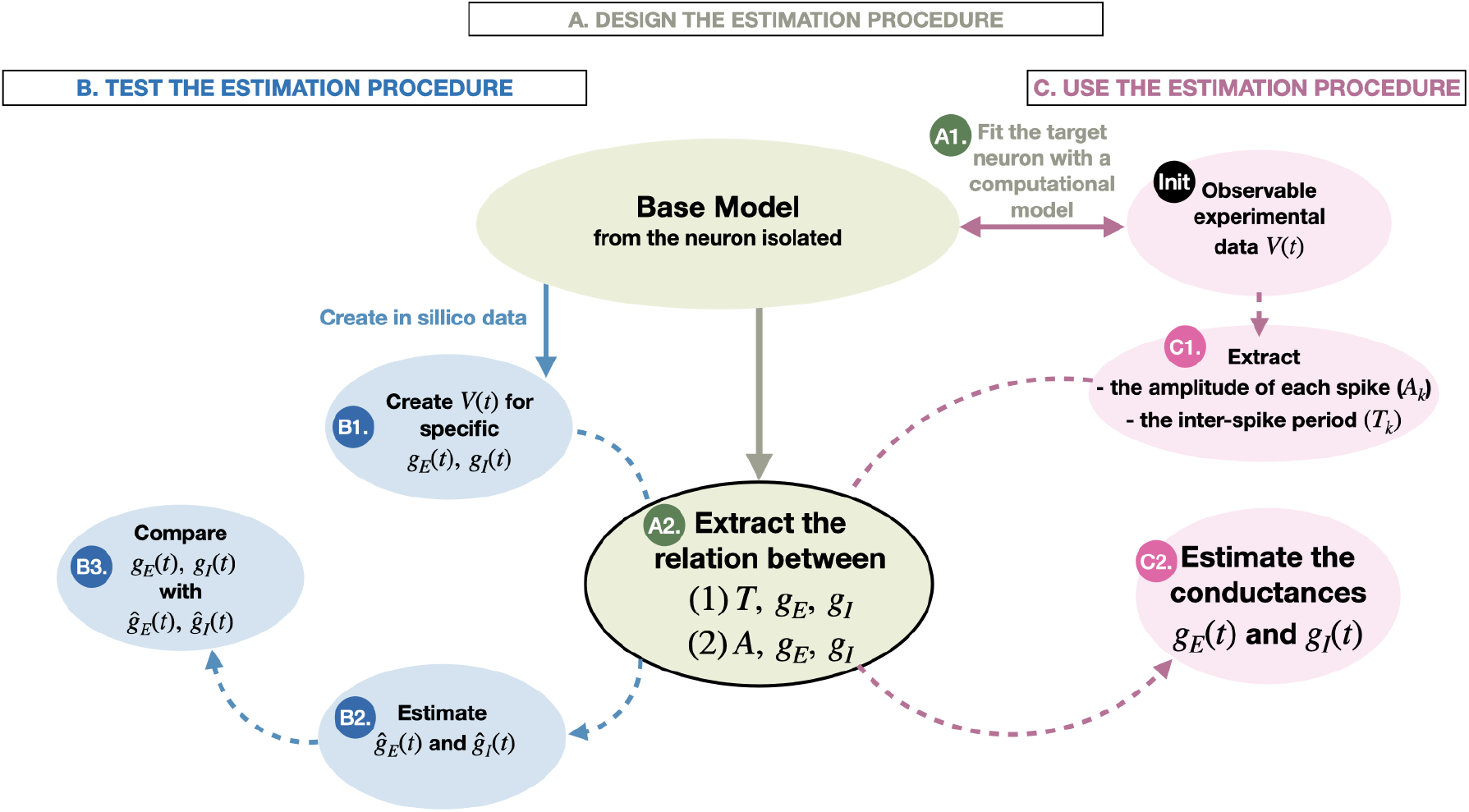
Schematic of the problem of synaptic conductance estimation. By considering a target neuron (step *Init*), its neuronal activity needs to be fitted to a base model in absence of synaptic currents impinging to the neuron (step *A1*). Then, the estimation procedure (**green pathway**) consists of, using the base model, extracting a relation between the conductances (*g*_*E*_, *g*_*I*_) and both the interspike period *T* and the amplitude *A* from the corresponding membrane potential *V* (*t*) calculated using the base model (step *A2*). From this relationship, the conductances can be estimated by providing both *A* and *T* of a particular membrane potential trace. To test the estimation procedure (**blue pathway**), we create *in sillico* membrane potential traces using the base model with prescribed representative synaptic conductance traces (step *B1*). Then, using the estimation procedure (step *A2*), we extract the conductances (step *B2*) and compare them to the original ones (step *B3*) to predict which are the errors caused by the procedure. Finally, the conductances impinging on the target neuron of the *Init* step can be estimated using the designed estimation procedure (**pink pathway**).

The article is organized as follows. Section 1 outlines the core estimation procedure (Subsections 1.1) and presents the specific neuronal model used as the base model for this framework (Subsection 1.2). Due to its relevance in E/I balance processes and its capability to generate membrane potential traces with a rich variety of amplitudes and frequencies in response to synaptic inputs, a stellate cell model is chosen to illustrate the procedure’s performance, although the method can be readily applied to other neural models. Later in this section, we introduce the different conductance traces used to test the method (Subsection 1.3). Section 2 presents and validates the method across different scenarios. Specifically, we first statistically demonstrate the accuracy of the estimation for constant synaptic conductance traces, followed by an analysis of how noise degrades performance by altering the neural dynamics and perturbing the relationship between (*T, A*) and (*g*_*E*_, *g*_*I*_) (Subsection 2.1). Next, we show how the estimation procedure accurately captures the time course of the inputs across a wide set of prescribed excitatory and inhibitory conductance profiles (Subsection 2.2). Finally, Section 3 concludes the paper with a brief discussion and concluding remarks.

## 1 Materials and methods

### 1.1 Estimation procedure

We consider a target neuron from which we want to estimate the excitatory and the inhibitory synaptic conductance that it is receiving per unit of time. For this purpose, a computational model that describes the membrane potential of the specific neuron when it is isolated or in absence of synaptic inputs is required. Assume this model being of the form

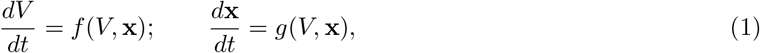

where *V* is the membrane potential and **x** other required variables like the gating variables. We add the synaptic current term *g*_*E*_(*t*)(*V* (*t*) − *V*_*E*_) + *g*_*I*_ (*t*)(*V* (*t*) − *V*_*I*_) to the membrane potential differential equation 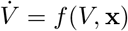 (the dot denotes derivative with respect to time), where *g*_*E*_ and *g*_*I*_ are the excitatory and the inhibitory conductance, and *V*_*E*_ and *V*_*I*_ are the excitatory and the inhibitory reversal potential, respectively.

Hence, with the new term, the equation for the membrane potential becomes 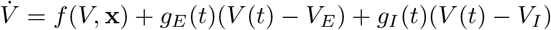 + *g*_*E*_(*t*)(*V* (*t*) − *V*_*E*_) + *g*_*I*_ (*t*)(*V* (*t*) − *V*_*I*_). We call this model the *base model*. Once the model is defined, we need to determine a realistic domain for the synaptic conductances; for simplicity, we assume this domain to be a rectangle ℛ = [*g*_*E,min*_, *g*_*E,max*_] × [*g*_*I,min*_, *g*_*I,max*_] in ℝ^2^. We denote by *S* the subset of ℛ where the steady-state solution of (1) is a limit cycle. We can also define functions *T* and *A* from *S* to ℝ assigning, respectively, the period and the amplitude of the limit cycle.

The estimation procedure is structured in several steps as follows. First, we sample a grid of constant synaptic conductances *G* = {(*g*_*E*_, *g*_*I*_)_*n*_ ∈ ℛ | *n* ∈ *N*}, where *N* is a discrete double-index set. We will typically choose *N* = [*N*_*E*_] × [*N*_*I*_], where [*n*] = {1, …, *n*}. For each element in *G*, we numerically integrate the system (1) and we compute the steady-state oscillation period, *T*, and the amplitude, *A*, of the membrane potential. This process yields two discrete tables, *T* = {*T* (*g*_*E*_, *g*_*I*_) | (*g*_*E*_, *g*_*I*_) ∈ *G*} and *A* = {*A*(*g*_*E*_, *g*_*I*_) | (*g*_*E*_, *g*_*I*_) ∈ *G*} In Figure 2 we show a continuous representation of *T* and *A* for the specific base model that will be introduced in Section 1.2.

**Fig 2.**
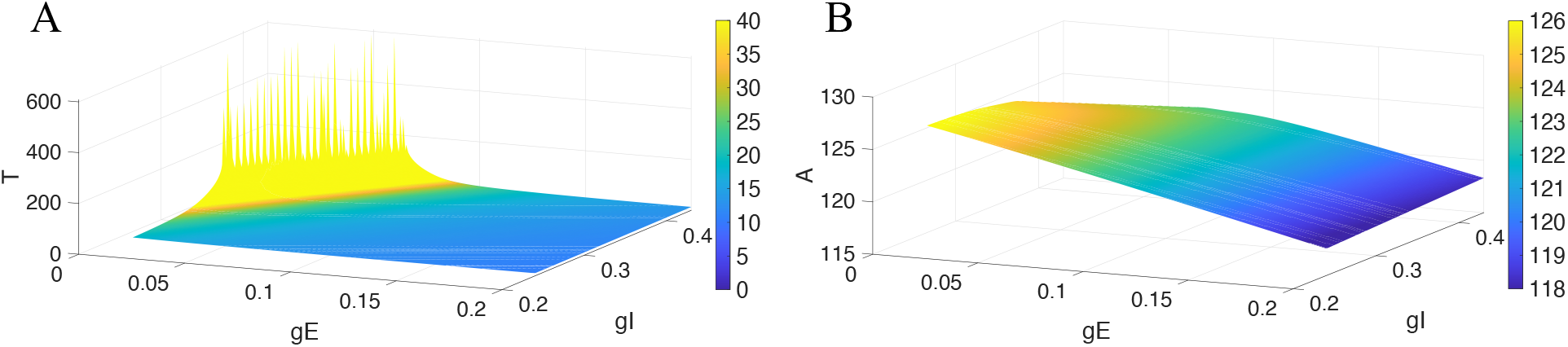
T and A functions. Considering the stellate cell model (see Section 1.2), Panel **A** and Panel **B** represent the spike period and the amplitude of oscillation obtained for different excitatory and inhibitory inputs, respectively. The period (in **A**) and the amplitude (in **B**) are also represented by the colorbars, which are capped at 40 *ms* and 126 *mV*, respectively.

Notice that, given a specific period *T*^∗^ and amplitude *A*^∗^, we can extract the closest pair (*g*_*E*_, *g*_*I*_) in *T* and *A* that provides *T*^∗^ and *A*^∗^. In order to obtain the full time course of the conductances, and based on the ideas in [25], we take advantage of the interspike intervals to compute local oscillatory periods and amplitudes. That is, given the membrane potential *V* (*t*) from which we want to extract the excitatory and inhibitory contribution, we compute the different interspike intervals and also the amplitude of each spike, obtaining, in this way, two respective sequences 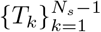 and 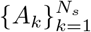, where *N*_*s*_ is the number of spikes in *V* (*t*). In order to have the same dimension for both sequences, we do not consider the amplitude of the first spike, and so the considered amplitude sequence is 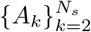, which we will recall as 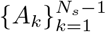 forconvenience.

Finally, per each pair (*T*_*k*_, *A*_*k*_), with *k* ∈ {1, …, *N*_*s*_ − 1}, and considering the discretized tables *T* and *A* obtained above, we look for the couple (*g*_*E*_, *g*_*I*_) that simultaneously minimizes the Euclidean distance between (*T* (*g*_*E*_, *g*_*I*_), *A*(*g*_*E*_, *g*_*I*_)) and (*T*_*k*_, *A*_*k*_), for (*g*_*E*_, *g*_*I*_) ∈ *G*. That is,

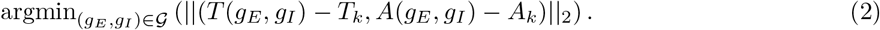

Let us call these pairs as (*gE*_*k*_, *gI*_*k*_), for *k* = 1, …, *N*_*s*_ − 1. Note that the minimization problem may yield multiple conductance pairs, leading to potential ambiguities in the estimation procedure. To break this tie, we always select the pair closest to the one obtained in the previous step.

#### 1.1.1 Improving the estimation procedure

Even though the latter procedure gives excellent estimations (see Section 2.1), the obtained pair (*gE*_*k*_, *gI*_*k*_) depends on the performed discretization of *g*_*E*_ and *g*_*I*_ considered to build up the tables *T* and *A*. To improve possible misestimations coming from this discretization, each pair (*gE*_*k*_, *gI*_*k*_) is readjusted by solving the 2-dimensional system

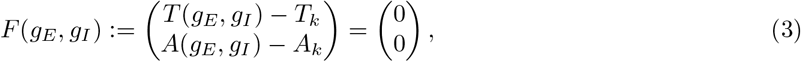

using a numerical version of the classical Newton’s method. More precisely, let us rewrite system (3) as *F* (*g*_*E*_, *g*_*I*_) = 0. Then, to apply the classic Newton’s method, we need *F* (*g*_*E*_, *g*_*I*_) being of class *C*^1^(*S*) with Jacobian matrix *DF* (*g*_*E*_, *g*_*I*_). Since *F* (*g*_*E*_, *g*_*I*_) = 0 has not an analytical expression, *DF* (*g*_*E*_, *g*_*I*_) is computed as the numerical gradient of the tables *T* and *A* defining the spacing between points in each direction as the discretization of *g*_*E*_ and *g*_*I*_. Subsequently, a two-dimensional cubic spline interpolation is applied to these surfaces to evaluate both *F* (*g*_*E*_, *g*_*I*_) and *DF* (*g*_*E*_, *g*_*I*_) as continuous functions during the iterative process. On the other hand, since the condition *C*^1^(*S*) is not assured, if the method does not converge for a specific pair of initial conditions, then we preserve the initial conditions as the solution of the system.

Ultimately, we obtain a sequence of synaptic conductance pairs 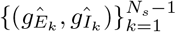. By selecting the *k*-th spike onset time, *t*_*k*_, as the corresponding onset time of the 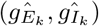 pair, we construct the two temporal sequences 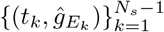 and 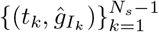, which are then interpolated to obtain the continuous time course of the excitatory and inhibitory conductances. In our simulations, we use the cubic spline method.

The estimation procedure is summarized in Algorithm 1 and visualized in Figure 3. The source code is publicly available at https://github.com/RosaDelic/ConductanceEstimation.git.

**Fig 3.**
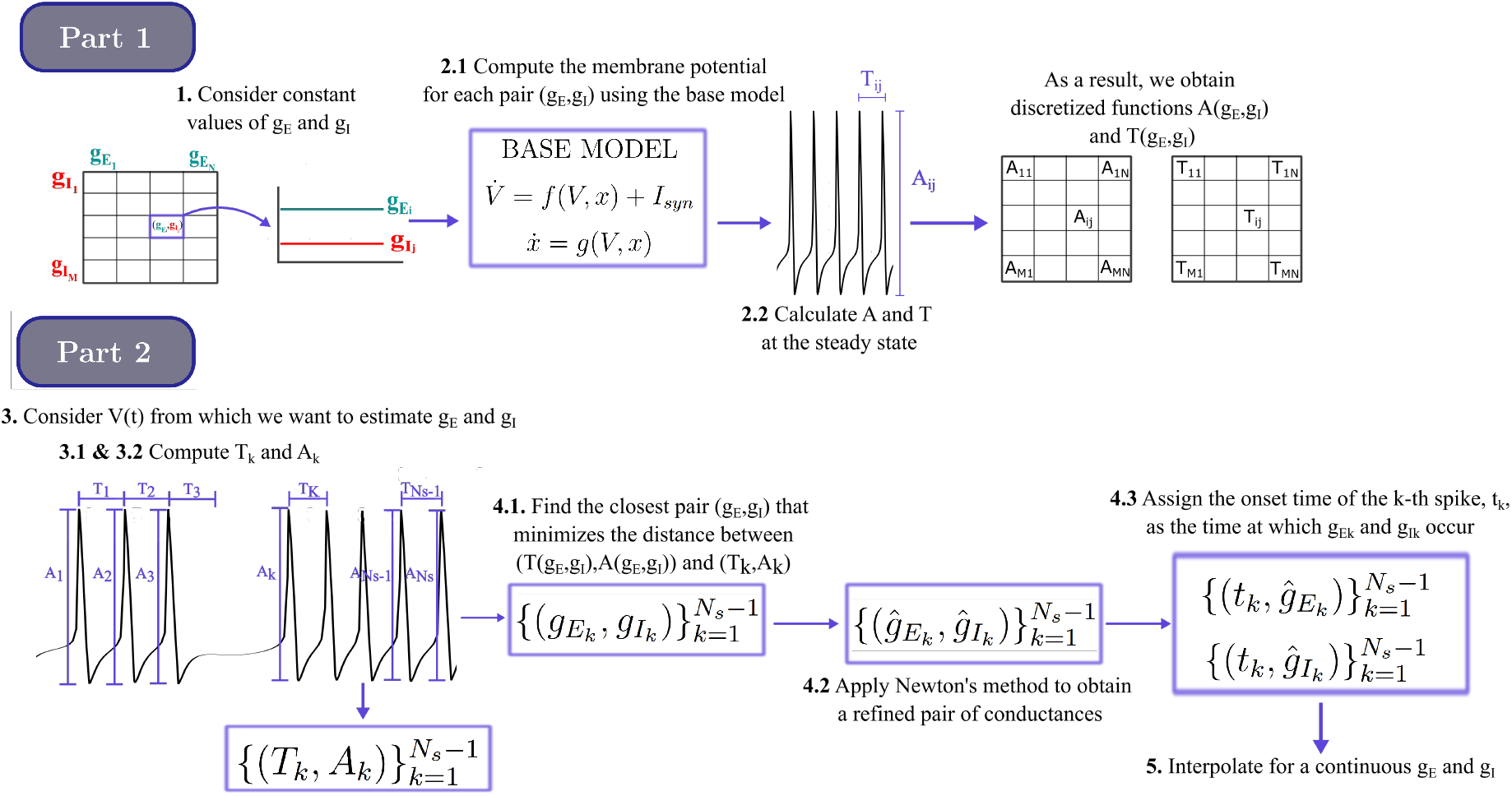
Representation of the estimation procedure described in Algorithm 1. Visualization of the different steps of the estimation procedure. Part 1 corresponds to the construction of tables *T* and *A* while Part 2 estimates the conductances from the given membrane potential. Steps are numerated as in Algorithm 1.

### 1.2 Neural model

As highlighted in the introduction, stellate cells play distinct functional roles within the navigational circuits of the medial entorhinal cortex (MEC) [28, 29] and the motor control networks of the cerebellum [30, 31]. In this paper, we consider the stellate cell model as our base model to exemplify the proposed estimation procedure, due to its critical involvement in E/I balance processes and its property of exhibiting different spike amplitudes depending on the strength of the external input. Nonetheless, our estimation procedure could be readily extended to any other neural model sharing these dynamic characteristics.

#### Algorithm 1

Estimation procedure

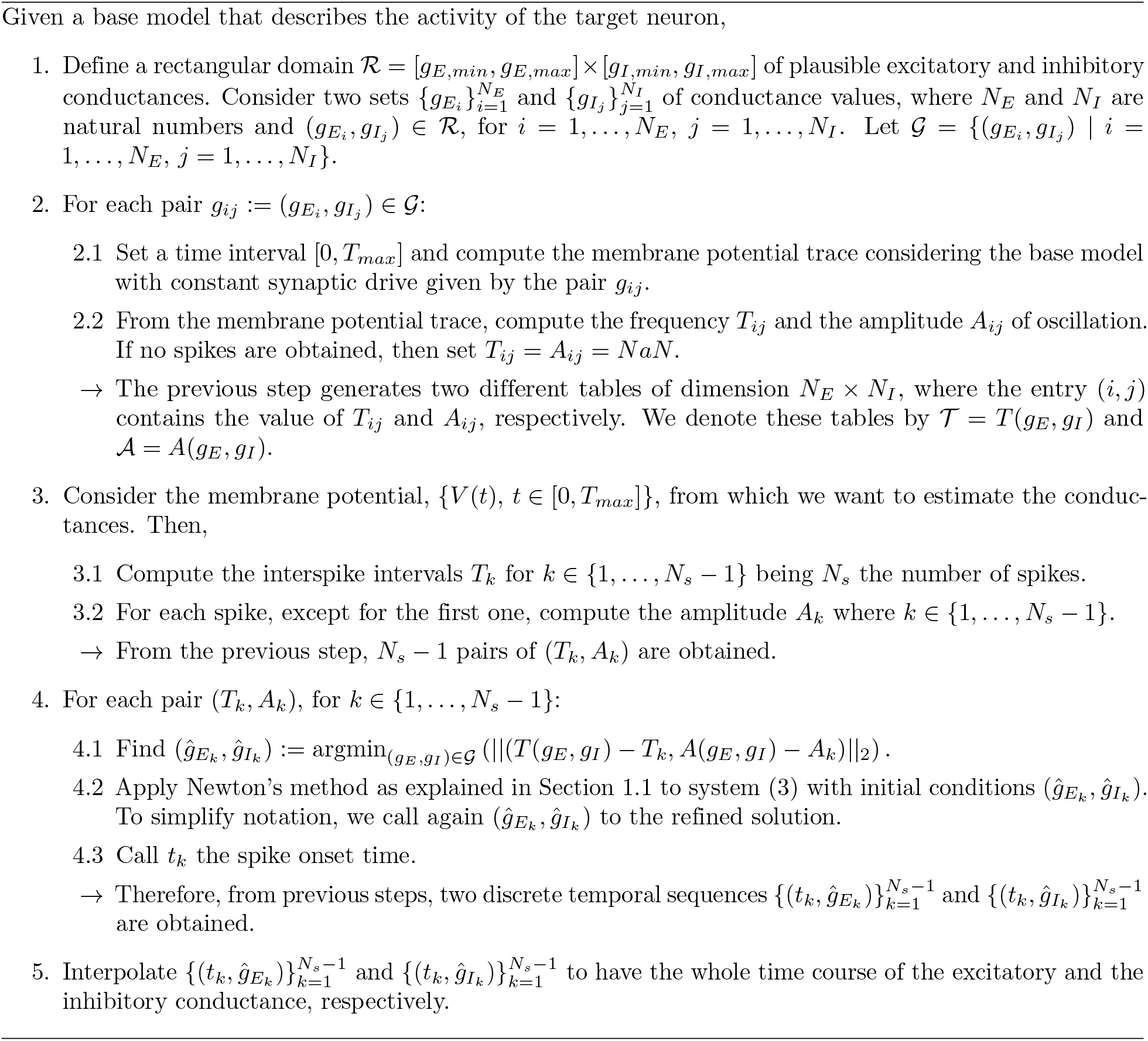

We consider the model proposed in [32], consisting of a spiking neuron driven by a sodium current (*I*_*Na*_), a delayed-rectifier potassium current (*I*_*K*_), a persistent sodium current (*I*_*NaP*_) and a hyperpolarization-activated current (*I*_*h*_). The dynamics of its membrane potential is described by

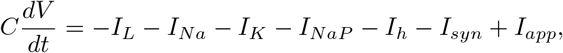

where *C* represents the membrane capacitance, *I*_*L*_ the leak current, *I*_*syn*_ the synaptic current, and *I*_*app*_ the applied current. All ionic currents are time- and voltage-dependent, described according to the Hodgkin-Huxley formalism (see S1 Appendix for full details on their formulation). The applied current *I*_*app*_ is supposed to be constant throughout the paper, and *I*_*syn*_ is detailed in the next subsection.

This model is characterized by the fact that both the spike amplitude and the period vary across different synaptic inputs (see Figure 4), providing the two observable quantities necessary to discern the respective contributions of excitatory and inhibitory currents.

**Fig 4.**
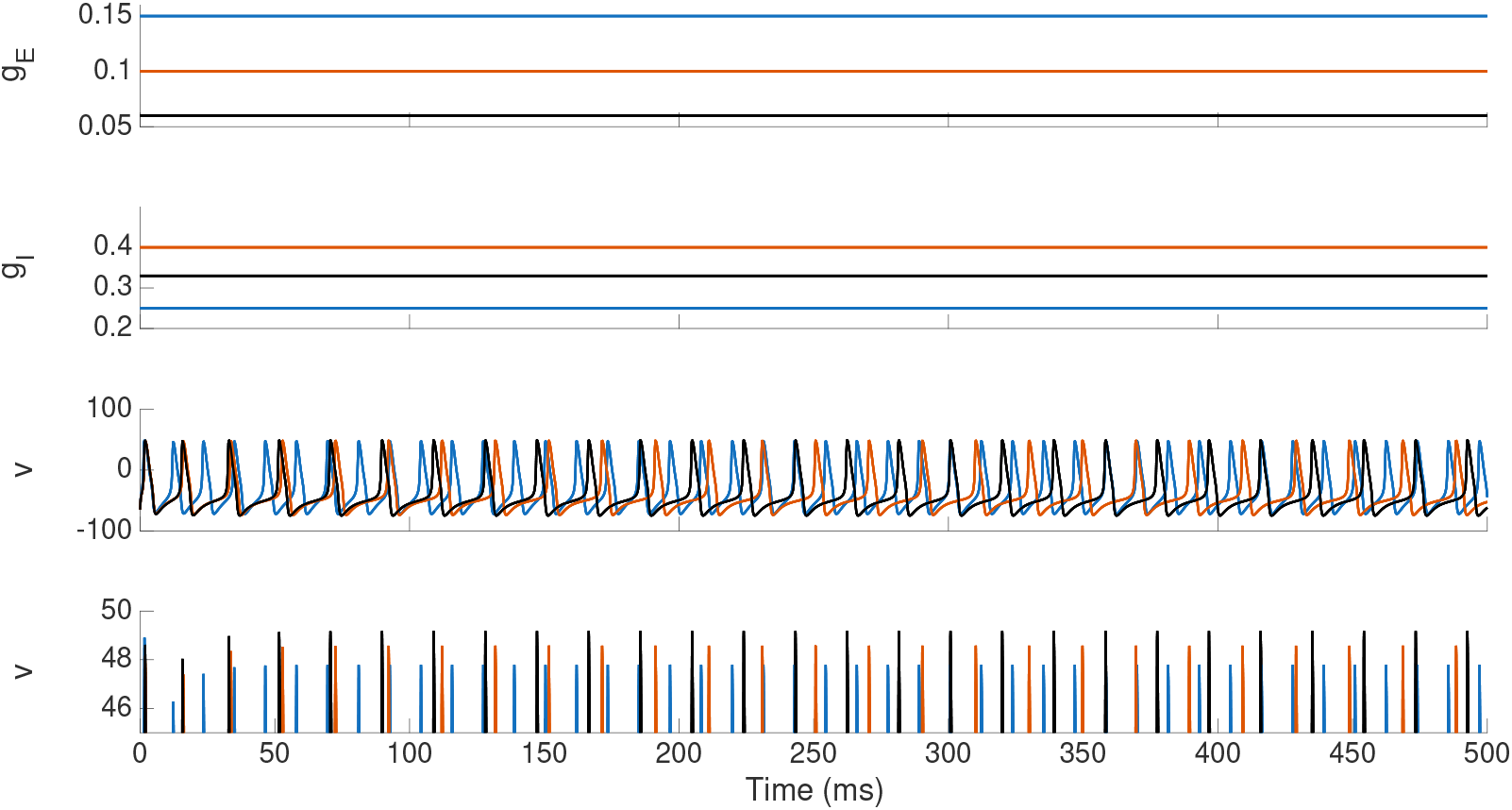
Example of membrane potential traces obtained from the stellate cell model using different conductance traces. From top to bottom, the different panels represent the excitatory conductance trace *g*_*E*_(*t*), the inhibitory conductance trace *g*_*I*_ (*t*), the resulting membrane potential trace *v*(*t*) generated by these synaptic inputs, and a zoomed-in view of *v*(*t*) showing spike peaks with different amplitudes. Different colors correspond to different constant conductances’ values: blue traces correspond to *g*_*E*_(*t*) = 0.15 and *g*_*I*_ (*t*) = 0.25; dark orange trace corresponds to *g*_*E*_(*t*) = 0.1 and *g*_*I*_ (*t*) = 0.4; and black trace corresponds to *g*_*E*_(*t*) = 0.06 and *g*_*I*_ (*t*) = 0.33.

### 1.3 Prescribed synaptic conductance traces

The synaptic drive impinging to a target neuron is formulated as

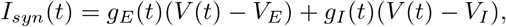

where *g*_*E*_(*t*) and *g*_*I*_ (*t*) refer to the excitatory and inhibitory conductances, respectively, while *V*_*E*_ and *V*_*I*_ are their reversal potentials, which are supposed to be constant (see S1 Appendix).

To assess the performance of the estimation procedure and to explore various scenarios of activity, we consider different prescribed conductance traces for *g*_*E*_ and *g*_*I*_, as explained below. Throughout this work, the conductance pairs used for the stellate cell model are restricted to the region

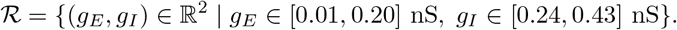

#### 1.3.1 Constant synaptic drive

In the constant case, we consider *g*_*E*_(*t*) = *g*_*E*_ and *g*_*I*_ (*t*) = *g*_*I*_. For statistical purposes, the (*g*_*E*_, *g*_*I*_) pairs are selected jointly over the two-dimensional domain ℛ using Latin Hypercube Sampling (LHS) to ensure a uniform coverage of the parameter space.

Although all obtained *in silico* traces show realistic spiking frequencies, not all combinations of constant *g*_*E*_ and *g*_*I*_ within this region guarantee that the target neuron exhibits the desired oscillatory behavior. Consequently, parameter sets yielding a non-spiking membrane potential trace are discarded (see red dots in Figure 5). We denote the set of all combined (*g*_*E*_, *g*_*I*_) values in ℛ that cause an oscillatory neural behavior by *S* (see blue dots in Figure 5).

**Fig 5.**
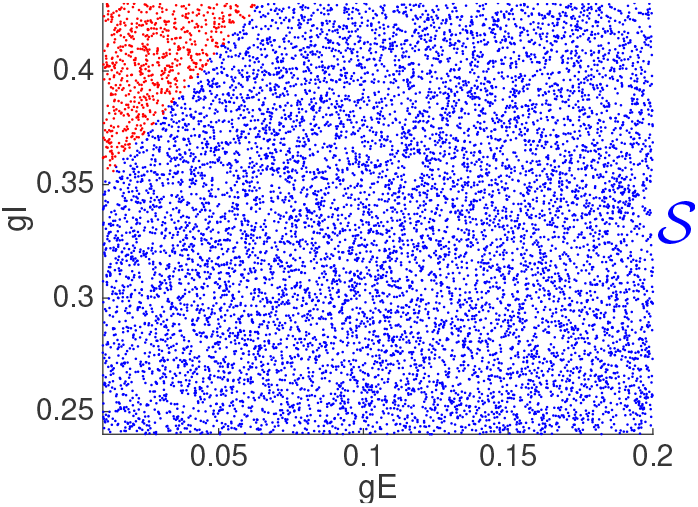
Pairs of (*gE, gI*) used in the Latin Hypercube Sampling (LHS). Each dot corresponds to a different pair of conductances used to generate different membrane potential traces. Blue dots represent the pairs of (*gE, gI*) that provide a spiking response on the membrane potential (a discrete subset in region *S*), while red dots correspond to those pairs causing a resting state response.

#### 1.3.2 Oscillatory synaptic drive with different phases

Testing the method across diverse E/I phase shifts—ranging from in-phase cortical balance to anti-phase motor rhythms and delayed feedback inhibition, ensures its ability to capture distinct, realistic physiological scenarios. To explore these cases, we consider excitatory and inhibitory oscillatory conductance traces that take into account phase-shifts. In particular,

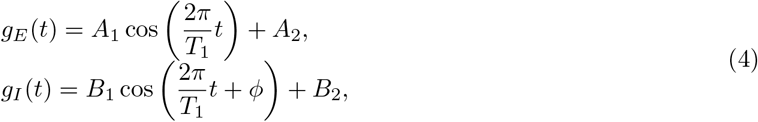

where the parameters are *T*_1_ = 300 *ms, A*_1_ = 0.055 *mV*, *A*_2_ = 0.105 *mV*, *B*_1_ = 0.075 *mV*, *B*_2_ = 0.335 *mV* and the phase *ϕ* belongs to [0, 2*π*).

#### 1.3.3 Double-frequency synaptic drive

To study how fast the conductances can change, we also consider *g*_*E*_(*t*) and *g*_*I*_ (*t*) having two different frequencies, one that varies more slowly than the spike frequency and another, with smaller amplitude but varying faster. That is,

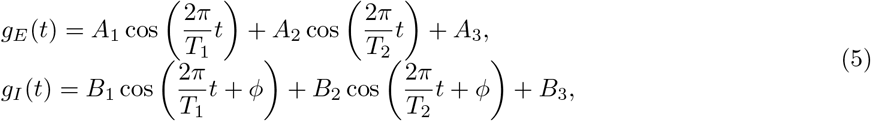

being *T*_1_ = 300 *ms, T*_2_ = 75 *ms, A*_1_ = *B*_1_ = 0.060 *mV*, *A*_2_ = *B*_2_ = 0.035 *mV*, *A*_3_ = 0.105 *mV*, *B*_3_ = 0.335 *mV*, and the phase *ϕ* belongs to [0, 2*π*).

#### 1.3.4 *In silico* synaptic drive

Finally, we use more realistic synaptic conductance traces extracted from a computational network, with a 1 *ms* resolution, that models the layer 4*Cα* of the primary visual cortex. See [33, 34] for more information on the data and Figure 10 for a visualization of the traces.

For simplicity, from now on we will denote *g*_*E*_ = *g*_*E*_(*t*) and *g*_*I*_ = *g*_*I*_ (*t*) if no confusion arises.

## 2 Results

We divide this section into two parts: first, in Section 2.1, we address the robustness of the proposed method, and second, in Section 2.2, we apply it to the selected model to evaluate the accuracy of the estimates.

### 2.1 The estimation procedure robustly estimates constant synaptic drives

The quality of the estimates obtained with our proposed method can be affected by several key factors: the intrinsic characteristics of the model, the choice of certain parameters in the numerical procedure, or the presence of noise. More precisely, we study: (1) the goodness of fit given by the accuracy of the computed functions *T* (*g*_*E*_, *g*_*I*_) and *A*(*g*_*E*_, *g*_*I*_), mainly determined by the choice of the discretization parameter *δ*; (2) possible improvements in the estimation after refining the results with Newton’s method (Step 4.2 in the Algorithm 1); and, (3) the quality of the estimations under different noise levels.

To perform the aforementioned examinations, we generate a set of 10^4^ different *in sillico* membrane potential traces, *V*_*j*_(*t*), with *j* ∈ {1, … 10^4^} and *t* ∈ [0, *T*_*max*_], using the stellate cell model described in Section 1.2. Each simulation has a different synaptic input, defined by a pair of constant excitatory and inhibitory conductances (*g*_*E*_, *g*_*I*_)_*j*_ ∈ *S* that have been randomly generated using the Latin Hypercube Sampling (LHS), which ensures a more uniform set of sample points across all possible values. Note that since the conductances are constant, the interspike intervals and amplitudes are constant along the time interval [0, *T*_*max*_]. Therefore, from Step 3 of Algorithm 1, we obtain a unique period-amplitude pair (*T*_*j*_, *A*_*j*_) for each *j* = 1, …, 10^4^. We will maintain these table of (*T*_*j*_, *A*_*j*_) values throughout Sections 2.1.1, 2.1.2, and 2.1.3, whereas the reference tables *T* and *A* will change depending on what is studied in each section.

#### 2.1.1 The accuracy of the estimation depends on the discretization

For simplicity, without loss of generality, we concentrate the discretization on a single parameter *δ*> 0. We define *N*_*E*_ and *N*_*I*_ in Step 1 of Algorithm 1 as *N*_*E*_ = ⌈(*g*_*E,max*_ − *g*_*E,min*_)/*δ*⌉ and *N*_*I*_ = ⌈(*g*_*I,max*_ − *g*_*I,min*_)/*δ*⌉; therefore, *δ* determines the grid spacing of the discrete set *G* and, consequently, the resolution of tables *T* and *A* defined in Step 2 of Algorithm 1 will depend on *δ*.

For each *δ* value and each *j* ∈ {1, …, 10^4^}, we apply Steps 4 and 5 of Algorithm 1 to (*T*_*j*_, *A*_*j*_) obtaining a single pair of estimated constant conductances (*ĝ*_*E,j*_, *ĝ*_*I,j*_). To evaluate the performance, we compute the mean absolute error (MAE) of the estimated conductances with respect to the corresponding point in the LHS. These averaged errors demonstrate that the estimation accuracy is determined by the choice of *δ*, remaining within the same order of magnitude (see Table 1).

**Table 1.** Errors obtained from the conductances discretization in *T* (*g*_*E*_, *g*_*I*_) and *A*(*g*_*E*_, *g*_*I*_) tables. Errors caused on the estimation of a constant synaptic conductance when different *g*_*E*_ and *g*_*I*_ discretizations (see first column) are considered in the estimation procedure in Algorithm 1. The second, third and fourth column show the averaged absolute error (MAE) of *g*_*E*_, *g*_*I*_ and the joined (*g*_*E*_, *g*_*I*_) pair, respectively, over 10^4^ different values of pairs of conductances. The last column shows the mean squared error (MSE) for the joined (*g*_*E*_, *g*_*I*_) case.

| $\delta$ | MAE ($g_E$) | MAE ($g_I$) | MAE ($g_E, g_I$) | MSE($g_E, g_I$) |
| --- | --- | --- | --- | --- |
| $5 \cdot 10^{-2}$ | $1.97 \cdot 10^{-2}$ | $3.45 \cdot 10^{-2}$ | $6.35 \cdot 10^{-2}$ | $1.18 \cdot 10^{-3}$ |
| $10^{-2}$ | $4.54 \cdot 10^{-3}$ | $7.93 \cdot 10^{-3}$ | $3.39 \cdot 10^{-2}$ | $1.09 \cdot 10^{-4}$ |
| $5 \cdot 10^{-3}$ | $2.50 \cdot 10^{-3}$ | $4.33 \cdot 10^{-3}$ | $2.99 \cdot 10^{-2}$ | $4.63 \cdot 10^{-5}$ |
| $10^{-3}$ | $5.47 \cdot 10^{-4}$ | $9.83 \cdot 10^{-4}$ | $2.63 \cdot 10^{-2}$ | $3.99 \cdot 10^{-6}$ |
| $5 \cdot 10^{-4}$ | $2.73 \cdot 10^{-4}$ | $5.37 \cdot 10^{-4}$ | $2.58 \cdot 10^{-2}$ | $1.11 \cdot 10^{-6}$ |

In particular, the estimation of both *g*_*E*_ and *g*_*I*_, separately, yields absolute errors of about half the discretization step *δ*, showing that, if the base model fits the neural activity, then the estimation is as accurate as the *T* (*g*_*E*_, *g*_*I*_) and *A*(*g*_*E*_, *g*_*I*_) functions. However, the absolute errors obtained for the estimated pair (*g*_*E*_, *g*_*I*_) are *O*(10^−2^) regardless of the value of *δ*. This fact is due to the difference in magnitude between *g*_*E*_ and *g*_*I*_ and also a compensation between the errors given in both cases separately. The smaller the value of *g*_*E*_, the larger the error in its estimation (see Figure 6A1), and the opposite occurs for *g*_*I*_ (the larger it is, the larger the error, see Figure 6A2). However, to account for the differences in magnitude of *g*_*E*_ and *g*_*I*_, we compute the mean squared error MSE obtained in the estimation of the joined (*g*_*E*_, *g*_*I*_) data, showing a considerable improvement in the estimation when *δ* decreases (see Table 1, last column).

**Fig 6.**
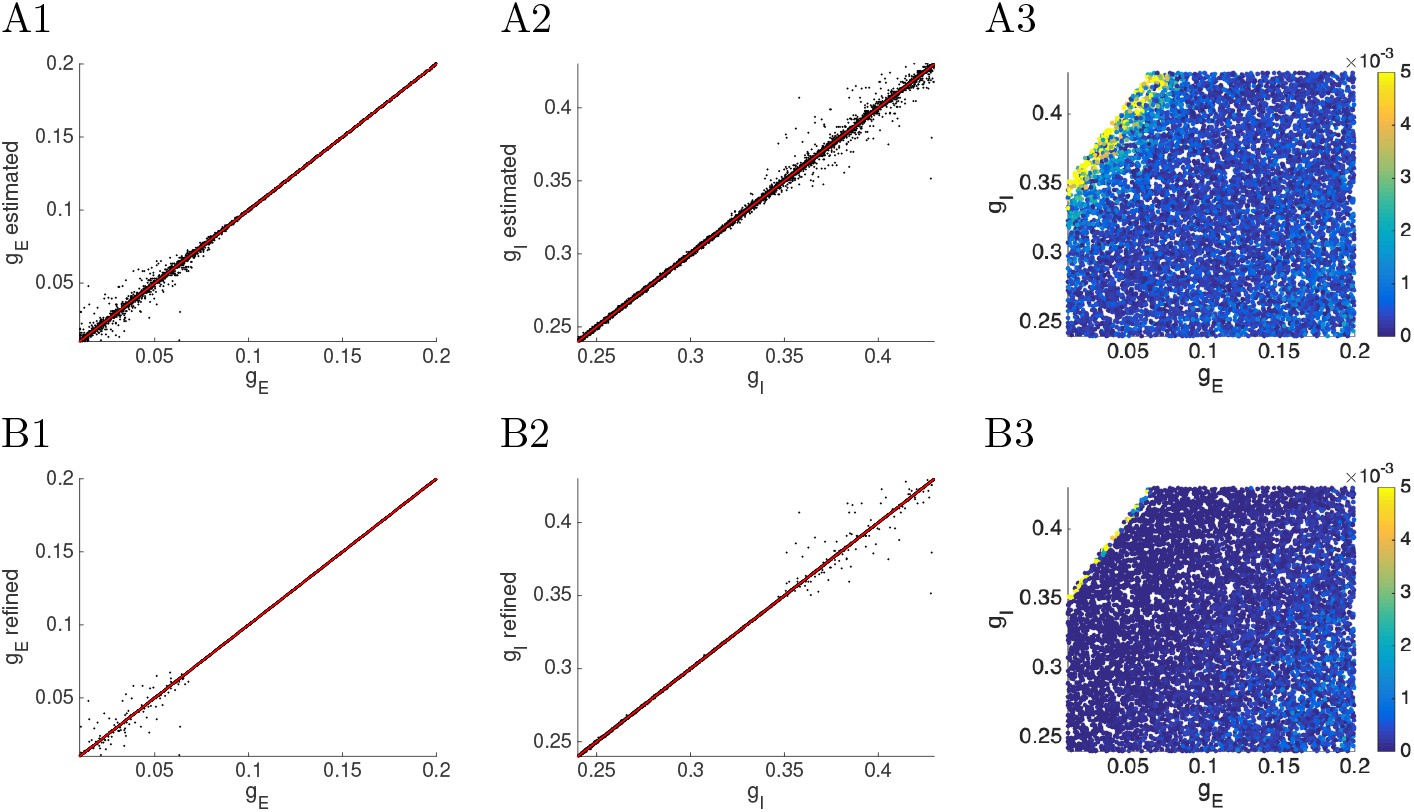
Estimated condutances using different constant conductance traces according to the LHS discretization. Panels A and B correspond to the estimation without and with applying Step 4.2 in Algorithm 1, respectively. First and second columns represent the scatter plots of the actual constant conductance versus the estimated ones, being the former column the excitation and the second the inhibition. In the four panels, the red line represents the identity line. At the third column, each dot describes the joined error of the estimated pair (*g*_*E*_, *g*_*I*_), computed using the Euclidian norm. In both panels A3 and B3, errors greater than 5· 10^−3^ have been cut for a better visualization.

By plotting the mean absolute errors for the (*g*_*E*_, *g*_*I*_) pairs in Figure 6 (panels A3 and B3), we observe that the errors increase as we approach the boundary of the oscillatory region *S*, while remaining significantly smaller in the rest of the domain. However, since in practice true conductances might be close to this boundary, these edge cases were intentionally kept in our analysis to provide a realistic assessment of the procedure’s performance.

#### 2.1.2 The estimation improves due to Newton’s refinement

By fixing *δ* = 10^−3^ and applying Step 4.2 of Algorithm 1, which implements Newton’s refinement explained in Section 1.1.1, the resulting conductance estimation errors are reduced compared to those obtained directly from the *T* and *A* tables with *δ* = 10^−3^ (see Table 2). Although the MSE decreases only slightly, both the minimum absolute error and the mean absolute error across realizations show substantial improvement.

**Table 2.** Errors obtained when estimating a constant synaptic conductance. Errors in the estimation procedure without (columns 2-4) and with (columns 5-7) applying Step 4.2 in Algorithm 1 with a discretization *δ* = 10^−3^. To compute these errors, we perform 10^4^ different estimations using a different pair (*g*_*E*_, *g*_*I*_) in each simulation. Second and fifth columns contain the minimum and maximum absolute value obtained across all the simulations. MAE columns represent the mean absolute error across these simulations and finally, MSE columns depict for the mean square error also across simulations. Euclidean norm has been used to compute the errors for the (*g*_*E*_, *g*_*I*_) pair.

|  | Errors from the Table search |  |  | Errors after applying Newton method |  |  |
| --- | --- | --- | --- | --- | --- | --- |
| | [min $E_a$, max $E_a$] | MAE | MSE | [min $E_a$, max $E_a$] | MAE | MSE |
| $g_E$ | $[1.33 \cdot 10^{-7}, 0.0521]$ | $5.47 \cdot 10^{-4}$ | $2.39 \cdot 10^{-6}$ | $[2.33 \cdot 10^{-9}, 0.0521]$ | $8.51 \cdot 10^{-6}$ | $1.14 \cdot 10^{-6}$ |
| $g_I$ | $[3.09 \cdot 10^{-8}, 0.0762]$ | $9.83 \cdot 10^{-4}$ | $5.59 \cdot 10^{-6}$ | $[2.86 \cdot 10^{-9}, 0.0762]$ | $2.70 \cdot 10^{-4}$ | $2.53 \cdot 10^{-6}$ |
| ($g_E, g_I$) | $[1.17 \cdot 10^{-5}, 0.433]$ | $2.63 \cdot 10^{-2}$ | $3.99 \cdot 10^{-6}$ | $[5.24 \cdot 10^{-8}, 0.433]$ | $2.94 \cdot 10^{-4}$ | $1.83 \cdot 10^{-6}$ |

This improvement is also reflected in Figure 6. As seen in Panels B1 and B2, fewer points lie outside the vicinity of the identity line compared to Panels A1 and A2, respectively. Moreover, concerning pair error (Panels A3 and B3), the errors in Panel B3 are significantly lower than those in Panel A3 for most conductance pairs. In fact, the points where the estimation does not improve as much are those that can be seen in the lower-right triangle, where *g*_*E*_ takes larger values while *g*_*I*_ takes smaller ones. Although some poorly estimated points remain near the non-spiking regime (upper-left), the estimation in this region is substantially improved compared to the results obtained before Newton’s refinement.

#### 2.1.3 Effects of the noise in the model

Estimating the conductances via Algorithm 1 requires either a model that perfectly fits the target neuron in the absence of synaptic inputs—as in the previous analysis—or the ability to extract the relationship of the excitatory and inhibitory conductances with their amplitude and oscillatory frequency or period from experiments. This relation can be obtained using firing-clamp techniques (see [26]). In this case, the data is subject to both measurement noise and trial-to-trial variability, which causes a loss of accuracy in the period and amplitude tables.

To assess the impact of these limitations, we add a zero-mean noise process to the membrane potential dynamics of the stellate cell model. Specifically, the neural dynamics are modeled as the stochastic differential equation

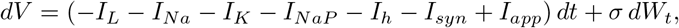

where *W*_*t*_ is a Wiener process, *σ* scales the noise and the rest of variables are fully described in Section 1.2 and 3.

By simulating this noisy stellate cell model across various values of *σ* and taking constant conductances from the set *G*, we obtain new *T* and *A* tables, which will be denoted by *T*_*σ*_ and *A*_*σ*_, respectively. Unlike the deterministic case with time-constant conductances, where a perfect oscillation is obtained, when noise is added, the oscillations become irregular. Thus, given a pair (*g*_*E*_, *g*_*I*_) ∈ *G*, we annotate the mean interspike intervals and the mean spike amplitude in the tables *T*_*σ*_ and *A*_*σ*_, respectively. The grid spacing in *G* is fixed to be *δ* = 10^−3^ for all *σ* values studied.

Our goal is to evaluate the quality of the conductance estimations as *σ* increases, using *T*_*σ*_ and *A*_*σ*_ as reference tables instead of *T* and *A* (corresponding to *T*_0_ and *A*_0_, respectively) used in the deterministic case.

Performing the estimation procedure (Steps 4 and 5 of Algorithm 1), we observe that, as expected, the estimation of excitatory, inhibitory, and paired conductances degrades as noise increases. Specifically, when transitioning from noise-free conditions (*σ* = 0) to a high-noise regime (*σ* = 1 *µA/*cm^2^), the mean absolute error (MAE) shifts by two orders of magnitude, from 10^−4^ to 10^−2^ (see Table 3), while the mean squared error (MSE) increases from 10^−6^ to 10^−4^. Given that the conductance values lie within ℛ, a maximum MAE on the order of 10^−2^ represents a minor deviation (≈ 2%) relative to their full scale, demonstrating that the method is robust to significant perturbations of the reference tables.

**Table 3.** Errors obtained in the estimation using tables. *T*σ and *Aσ*, which have been constructed from membrane potentials generated with different noise levels driven by conductances in *G*. Here, we fix the grid spacing of the discrete set *G* to *δ* = 10^−3^. Newton’s refinement has not been applied in these estimations.

| $\sigma$ | MAE ($g_E$) | MAE ($g_I$) | MAE ($g_E, g_I$) | MSE ($g_E, g_I$) |
| --- | --- | --- | --- | --- |
| 0 | $5.47 \cdot 10^{-4}$ | $9.83 \cdot 10^{-4}$ | $2.63 \cdot 10^{-2}$ | $3.99 \cdot 10^{-6}$ |
| $10^{-2}$ | $6.13 \cdot 10^{-3}$ | $9.58 \cdot 10^{-4}$ | $3.59 \cdot 10^{-2}$ | $6.99 \cdot 10^{-5}$ |
| $10^{-1}$ | $8.05 \cdot 10^{-3}$ | $1.26 \cdot 10^{-2}$ | $3.93 \cdot 10^{-2}$ | $1.21 \cdot 10^{-4}$ |
| 1 | $1.02 \cdot 10^{-2}$ | $2.17 \cdot 10^{-2}$ | $4.85 \cdot 10^{-2}$ | $4.51 \cdot 10^{-4}$ |

### 2.2 The estimation procedure captures the temporal evolution of the synaptic conductances

Although good results are obtained when constant values of conductances are estimated, conductances are time-varying. As shown in [25, 27], a good strategy to address this problem is to consider the inter-spike intervals (ISIs) as a time period where the conductances are constant and thus be able to apply the estimation procedure on the different ISIs to obtain the time course of the conductances. However, if the conductances vary faster than the time between spikes, the intermediate conductance values will not be properly estimated.

In this section, we explore the goodness of the estimation method across different scenarios of time-varying synaptic conductances. In Section 2.2.1, we consider different oscillation frequencies, both with and without phase shifts between the excitatory and inhibitory contributions, to explore a wide range of temporal relationships between *g*_*E*_(*t*) and *g*_*I*_ (*t*) (see the synaptic drive descriptions in Sections 1.3.2 and 1.3.3). We also examine the impact of measurement noise within this framework. Finally, in Section 2.2.2, we present the estimation results for a more irregular trace obtained from *in silico* data (Section 1.3.4).

To conduct this study, we generate the membrane potential using the stellate cell model (see Section 1.2) for different conductance traces. Then, treating the conductances as unknown, we estimate them using Algorithm 1 with the stellate cell model as the base model, followed by Newton’s refinement (with *δ* = 10^−3^). Finally, the estimated conductances are compared to the actual ones to see the accuracy of the estimation.

#### 2.2.1 Oscillatory synaptic drive

##### Phase-shift effects

To explore a comprehensive range of physiological conditions, we test our procedure across different oscillatory conductance traces for the synaptic drive (see equations (4) and (5)) with various phase shifts between *g*_*E*_(*t*) and *g*_*I*_ (*t*), specifically examining in-phase, anti-phase, and intermediate quadrature configurations. While an in-phase relationship reflects a tight, co-tuned E/I balance common in cortical networks, an anti-phase configuration represents an alternating computational mechanism typical of rhythmic motor circuits [35]. Furthermore, intermediate phase shifts, simulate biologically realistic scenarios of feed-forward or feedback inhibition, where a strict temporal delay exists between excitation and its counteracting inhibitory response [36]. Testing this full spectrum of phase relationships ensures that our estimation method is versatile enough to capture distinct operational states of the neuron, regardless of whether the synaptic inputs are synchronized, delayed, or mutually exclusive.

In Figure 7A,B we show how both the excitatory and the inhibitory conductance traces are well estimated when both conductances are in-phase (*ϕ* = 0). Although small errors are observed in the estimation of the inhibitory conductance trace when it becomes more irregular (double-frequency cases, see panel B, 4th column), they do not affect the reconstructed membrane potential (panel B, 3rd column). Indeed, both potential traces virtually overlap, without altering its frequency or its amplitude. When considering a single oscillation frequency, the MSEs obtained in these cases are on the order of 10^−5^ or even lower, whereas they are on the order of 10^−4^ when two frequencies are considered (see Table 4, first row). We emphasize that these MSEs have been calculated considering the estimated conductances after interpolating the results using a cubic spline interpolation. Errors prior to interpolation are reduced, in most cases, by one order of magnitude.

**Table 4.**
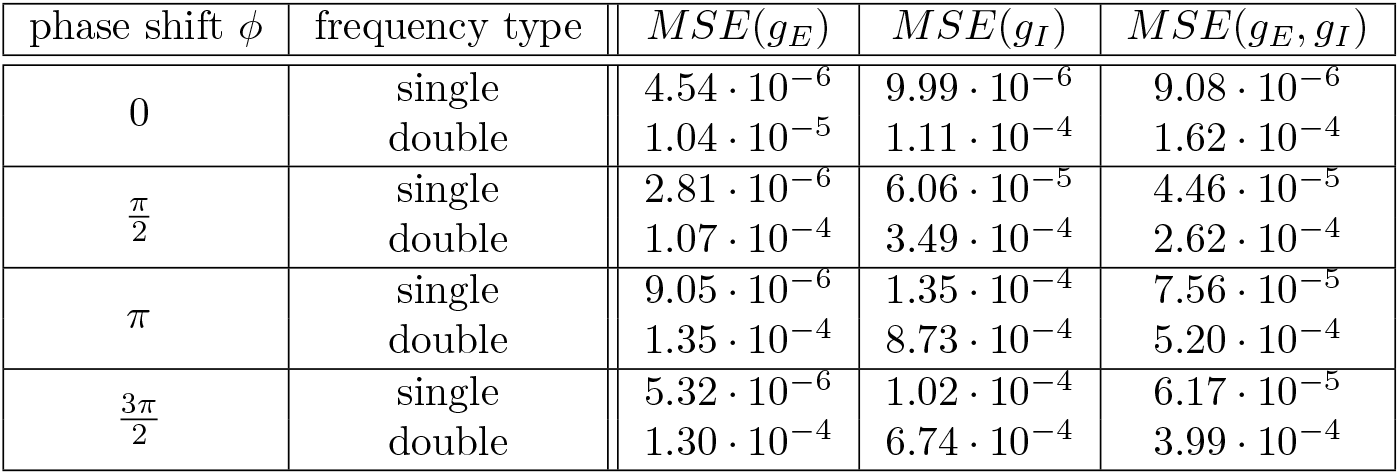
Estimation errors under oscillatory conductance inputs: effect of phase-shift and the number of frequencies of the synaptic input. Mean squared errors of the excitatory (3rd column), inhibitory (4th column) and the join error (5th column) are shown. First column contains the phase-shift between the excitatory and the inhibitory conductance traces. Second column indicates whether the oscillatory inputs have one or two frequencies. Error values have been obtained by comparing the actual and the estimated conductance traces considering a discretization parameter *δ* = 10^−3^ and the estimation after performing the Newton refinement.

| phase shift $\phi$ | frequency type | $MSE(g_E)$ | $MSE(g_I)$ | $MSE(g_E, g_I)$ |
| --- | --- | --- | --- | --- |
| 0 | single | $4.54 \cdot 10^{-6}$ | $9.99 \cdot 10^{-6}$ | $9.08 \cdot 10^{-6}$ |
| | double | $1.04 \cdot 10^{-5}$ | $1.11 \cdot 10^{-4}$ | $1.62 \cdot 10^{-4}$ |
| $\frac{\pi}{2}$ | single | $2.81 \cdot 10^{-6}$ | $6.06 \cdot 10^{-5}$ | $4.46 \cdot 10^{-5}$ |
| | double | $1.07 \cdot 10^{-4}$ | $3.49 \cdot 10^{-4}$ | $2.62 \cdot 10^{-4}$ |
| $\pi$ | single | $9.05 \cdot 10^{-6}$ | $1.35 \cdot 10^{-4}$ | $7.56 \cdot 10^{-5}$ |
| | double | $1.35 \cdot 10^{-4}$ | $8.73 \cdot 10^{-4}$ | $5.20 \cdot 10^{-4}$ |
| $\frac{3\pi}{2}$ | single | $5.32 \cdot 10^{-6}$ | $1.02 \cdot 10^{-4}$ | $6.17 \cdot 10^{-5}$ |
| | double | $1.30 \cdot 10^{-4}$ | $6.74 \cdot 10^{-4}$ | $3.99 \cdot 10^{-4}$ |

**Fig 7.**
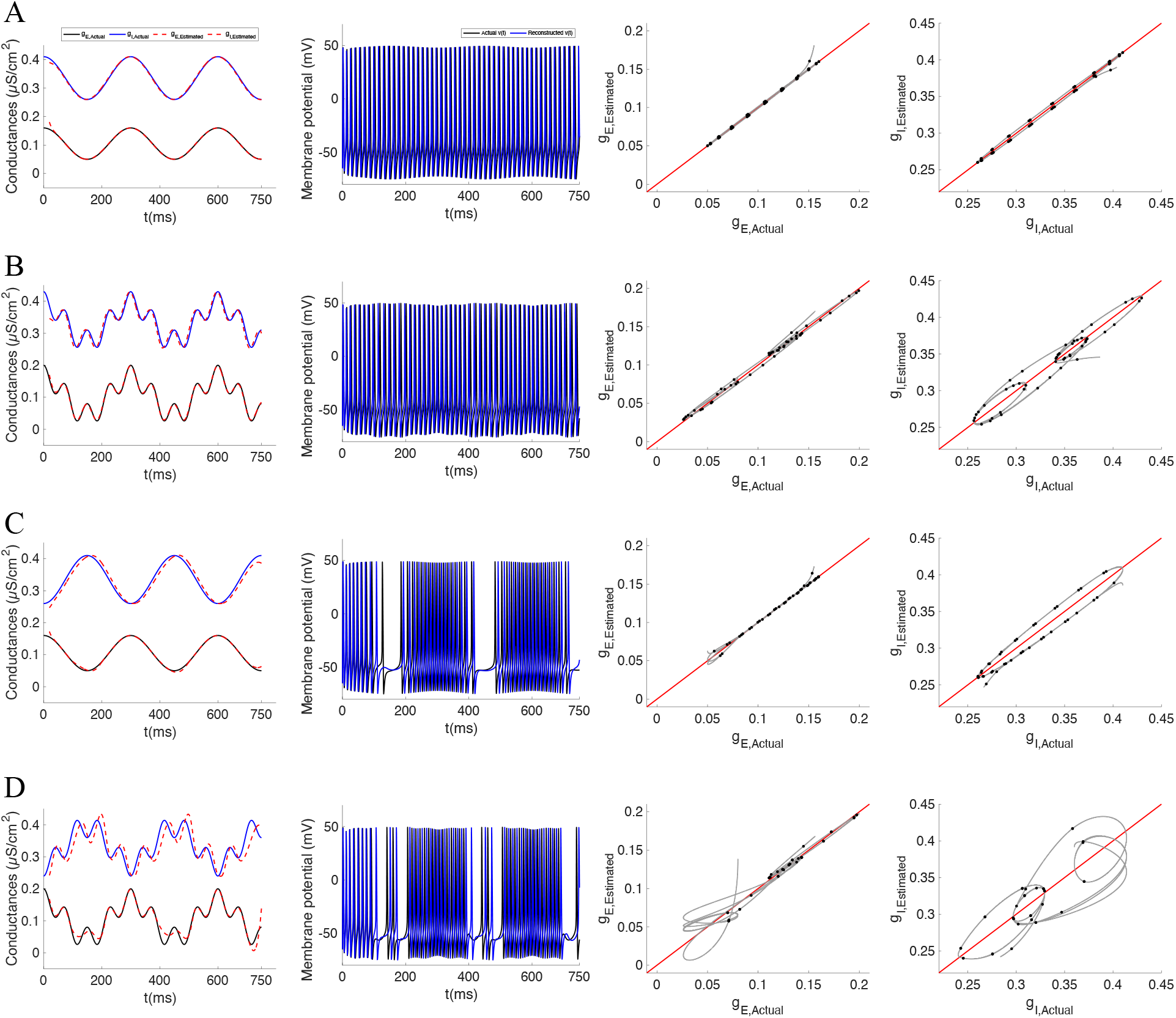
Estimation of the conductances time course for different oscillation frequencies and phases. Rows A and B show, for different oscillation frequencies respectively, the results when the excitatory and the inhibitory conductances are in-phase (*ϕ* = 0). Rows C and D correspond to the same oscillation frequencies as A and B respectively, but when the excitatory and the inhibitory conductances are anti-phase (*ϕ* = *π*). The equations and rest of parameters for the conductances in rows A and C are provided in equation (4) while those for rows B and D are provided in equation (5). From left to right, the different columns depict for: (1) the actual conductance traces (excitatory conductance in black and the inhibitory one in blue) and the corresponding estimated conductances provided by the estimation procedure (dashed red lines); (2) the actual membrane potential trace (black trace) and the reconstructed one by considering the estimated conductances (blue trace); (3) the actual excitatory conductances trace versus the estimated ones; and (4) the actual inhibitory conductances versus the estimated ones. In the 3th and 4th columns, black dots are the considered values in the estimation while the grey trace represents the interpolation of these values (the traces shown in the panel of corresponding 1st column). In all panels of the 3rd and 4th columns, the red line represents the identity line, the dots correspond to the discretized estimated conductances while the black traces show the conductances after interpolate them using the cubic spline interpolation.

In order to explore the out-of-phase configurations, we consider three different phase shifts, *ϕ*, such that 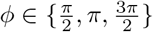. In Figure 7C,D we present the anti-phase case (*ϕ* = *π*), where a slight loss of precision in the estimation of the conductances can be appreciated. 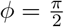 and 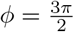 cases are shown in the S1 Fig, where similar results are present. To quantify this loss of precision, we compute the MSE values (see Table 4). Although the overall order of magnitude remains invariant, the error coefficient increases as *ϕ* approaches the anti-phase state (see also Figure 8, red traces in panels A.b,d and B.b,d, for a comprehensive evaluation across the full range of phase shifts). However, even though we obtain more inaccurate results as *g*_*E*_ and *g*_*I*_ become more out-of-phase, both figures show that these inaccuracies do not substantially affect the reconstructed membrane potential.

**Fig 8.**
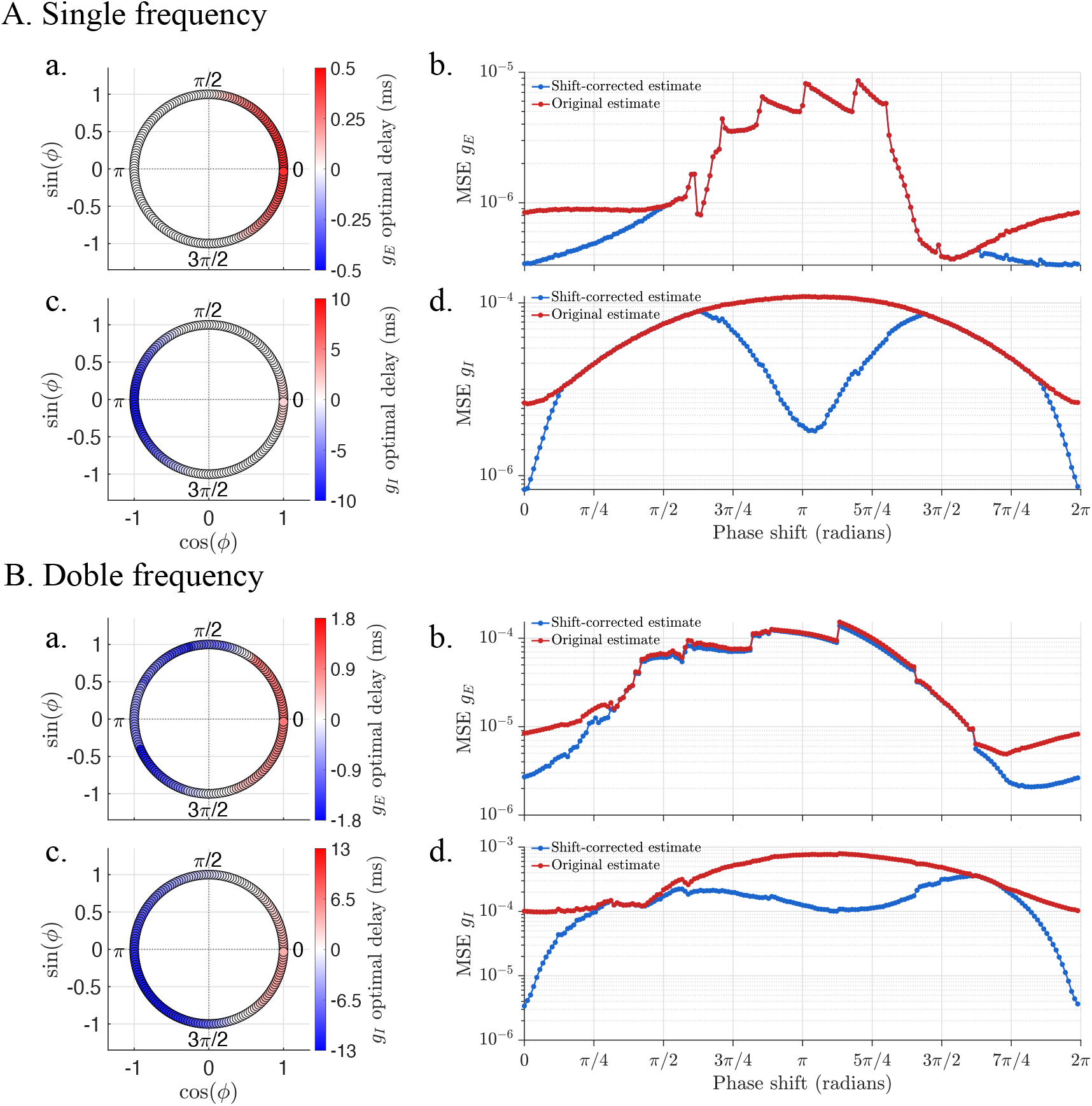
Effect of phase shift between *g*_*E*_(*t*) and *g*_*I*_ (*t*) on oscillatory synaptic conductance estimation. Panels A and B show the results when the excitatory and the inhibitory conductances are composed of a single or two oscillatory frequency components, respectively. At each panel, first column depicts the optimal temporal offsets, *τ*_opt,*E*_ (subpanels A.a and B.a) and *τ*_opt,*I*_ (subpanels A.c and B.c, that maximizes the correlation between the estimated and original excitatory (*g*_*E*_) and inhibitory (*g*_*I*_) synaptic conductances as a function of the imposed phase shift. Here, positive (red) and negative (blue) values indicate that the estimated conductance must be advanced or delayed, respectively, to achieve the best alignment. The second column represents the mean squared error (MSE) of the *g*_*E*_ (subpanels A.b and B.b) and *g*_*I*_ (subpanels A.d and B.d) reconstructions, before (original estimate, in red) and after (shift-corrected estimate, in blue) temporal alignment using the optimal offset shown in the first column. In all panels, the phase shifts *ϕ* are evaluated across the full range [0, 2*π*).

Figure 7 and S1 Fig suggest that the lower estimation performance observed under certain oscillatory regimes is not caused by a failure to capture the overall morphological waveform or amplitude of the inputs, but rather by a systematic horizontal time lag affecting both conductances. To correct this temporal displacement, we implemented a cross-correlation analysis between the true and the estimated signals as a post-processing step. This analysis covers both single and double oscillatory frequencies across a wide range of phase values, with *ϕ* ∈ [0, 2*π*). Specifically, the optimal time shift, *τ*_opt,*x*_, for *x* ∈ {*E, I*}, is determined by finding the argument that maximizes the cross-correlation function such that

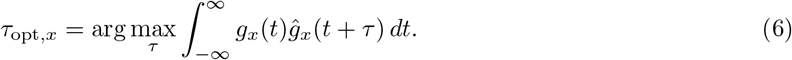

As illustrated in Figure 8, by shifting the estimated traces horizontally by *τ*_opt,*x*_, the realigned conductance *ĝ*_*x*,aligned_(*t*) = *ĝ*_*x*_(*t* + *τ*_opt,*x*_) exhibits a remarkable alignment with the true input.

The effect of phase shifts on conductance reconstruction is first analyzed for the single-frequency case in Figure 8A. For the excitatory conductance, positive values of *τ*_opt,*E*_, corresponding to an advance of the estimated conductance, are observed for phase shifts between 0 and *π/*2 and between 3*π/*2 and 2*π*, whereas no shift is observed for intermediate phases (subpanel A.a). This correction substantially reduces the reconstruction error (subpanel A.b): the original estimate exhibits MSE values of approximately 10^−5^ for phase shifts *ϕ* ∈ (*π/*2, 3*π/*2), and around 10^−6^ elsewhere. Applying *τ*_opt,*E*_ further decreases the MSE, particularly in the latter intervals.

A similar phase dependence is observed for the inhibitory conductance (Figure 8, subpanels A.c and A.d), although with an opposite correlation pattern. The optimal shift map displays negative cross-correlation values for *ϕ* ∈ (*π/*2, 3*π/*2) (subpanel A.c). Before correction, the *g*_*I*_ reconstruction error increases from 10^−5^ to a maximum of 10^−4^ near *π* and then decreases towards 2*π* (subpanel A.d). After applying *τ*_opt,*I*_, the MSE is markedly reduced around *π*, producing a local minimum of less than 10^−5^ and two maxima near *π/*2 and 3*π/*2. These results confirm that temporal misalignment is a major source of reconstruction error for both conductances.

Next, in Figure 8B, we examine how these observations change when the synaptic conductances contain two oscillatory frequency components. In contrast to the single-frequency case, optimal shift maps for both *g*_*E*_ and *g*_*I*_ (subpanels B.a and B.c) exhibit predominantly negative cross-correlation values over a broad phase interval extending approximately from *π/*2 to 3*π/*2. For the excitatory conductance, small regions of positive phase shift are observed in the first and fourth quadrants (subpanel B.a). A similar pattern is observed for the inhibitory conductance (subpanel B.c), although the region of positive phase shift is considerably narrower and mainly confined to phases close to 0. Moreover, the magnitude of the phase shift notably increases. The maximum errors increase by one order of magnitude with respect to the single-frequency case. After the phase-shift correction (blue curves in subpanels B.b and B.d) the improvement of the MSEs follows a similar pattern to the single-frequency case.

##### Noise effects

When considering noise in the stellate cell model, the time course of the synaptic conductances and their oscillatory behavior remains well predicted (see Figure 9, last two rows). However, the MSE across different phase shifts increases consistently with the noise level, reaching values on the order of 10^−3^ for higher noise regimes (*σ* = 1; see Figure 9, first two rows). Note that, similar to the double-frequency case (Figure 8B), at a low noise level (*σ* = 0.01), the MSE values are higher when approaching the quadrature phases (*ϕ* = *π/*2 and *ϕ* = 3*π/*2; see the magenta curves in the first two rows of Figure 9). Moreover, S2 Fig, corresponding to low level of noise (*σ* = 0.01), shows that the improvement when applying the phase-shift correction is qualitatively similar to the noise-free case (Figure 8). However, when noise increases substantially, for instance up to *σ* = 0.1 (see S3 Fig), the phase-shift correction loses effectiveness.

**Fig 9.**
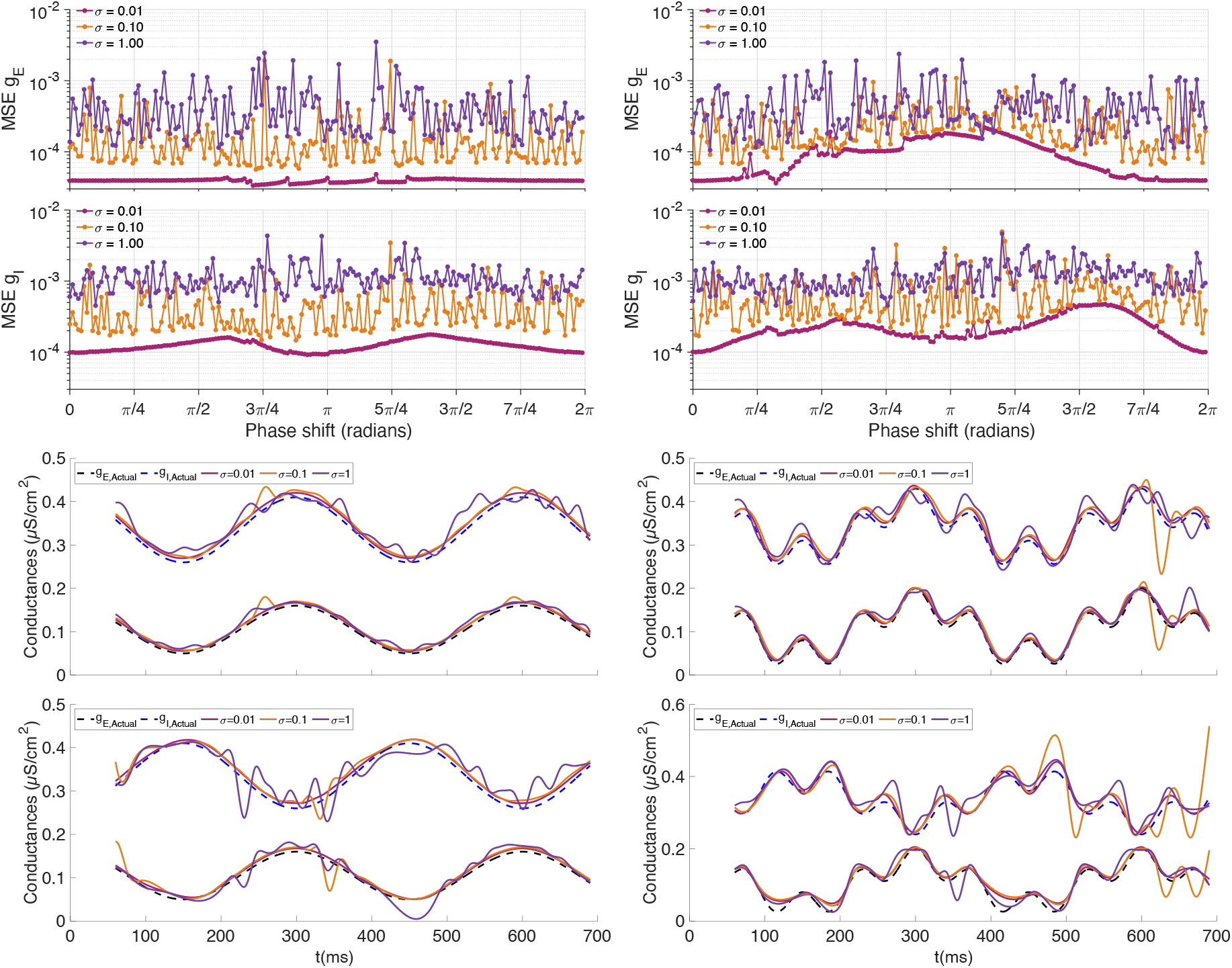
Estimation of the conductance time courses under different noise magnitudes. All panels display the estimation performance across various noise levels introduced into the membrane potential equation of the baseline system (noisy stellate model) prior to extracting the *T* _*σ*_ and *A* _*σ*_ tables, specifically for *σ* = 0.01 (magenta traces), *σ* = 0.1 (orange traces), and *σ* = 1 (purple traces). The left columns showcase the performance when single-frequency oscillatory conductance traces are considered, whereas the right columns correspond to double-frequency oscillations. From top to bottom, the rows depict: (1) the mean squared error (MSE) for the excitatory conductances, (2) the MSE for the inhibitory conductances, (3) an illustrative example of the time-course estimation for *g*_*E*_ (black) and *g*_*I*_ (blue) in-phase, and (4) the same for the out-phase case. In rows (3) and (4), the true inputs are represented by dashed lines (black for excitatory and blue for inhibitory), while the estimated traces are shown in the colors corresponding to each noise level *σ*.

#### 2.2.2 In sillico synaptic drive

Finally, we explore the efficacy of the procedure considering more realistic synaptic conductance traces, which have been obtained *in sillico* from a network that models the layer 4*Cα* of the primary visual cortex of a cat (see Section 1.3.4).·In this case, the inhibitory conductance abruptly changes in time (faster than the spiking frequency). Because of that, the estimation procedure is only able to capture the main oscillation of both *g*_*E*_ and *g*_*I*_ (see Figure 10A), presenting higher errors in the inhibitory case such that *MSE*(*g*_*I*_) = 2.38·10^−4^ versus *MSE*(*g*_*E*_) = 2.78·10^−6^. On the other hand, considering the mean conductances over time, we also obtain that the excitatory mean conductance is well estimated, being the actual mean *g*_*E*_ equal to 0.0746 and the estimated one equal to 0.0762 (absolute errors *O*(10^−3^)). However, the mean inhibitory conductance error is of order *O*(10^−2^), being the actual mean *g*_*I*_ equal to 0.353 and the estimated one equal to 0.337. These errors are reflected in the reconstructed membrane potential, see Figure 10B, where it can be appreciated that the poor estimation of inhibitory conductance oscillations ends up causing the reconstructed voltage to have periods of fewer spikes than in the original neuronal activity, followed by periods of higher oscillation frequency than it should be (see traces after the 400 *ms*).

**Fig 10.**
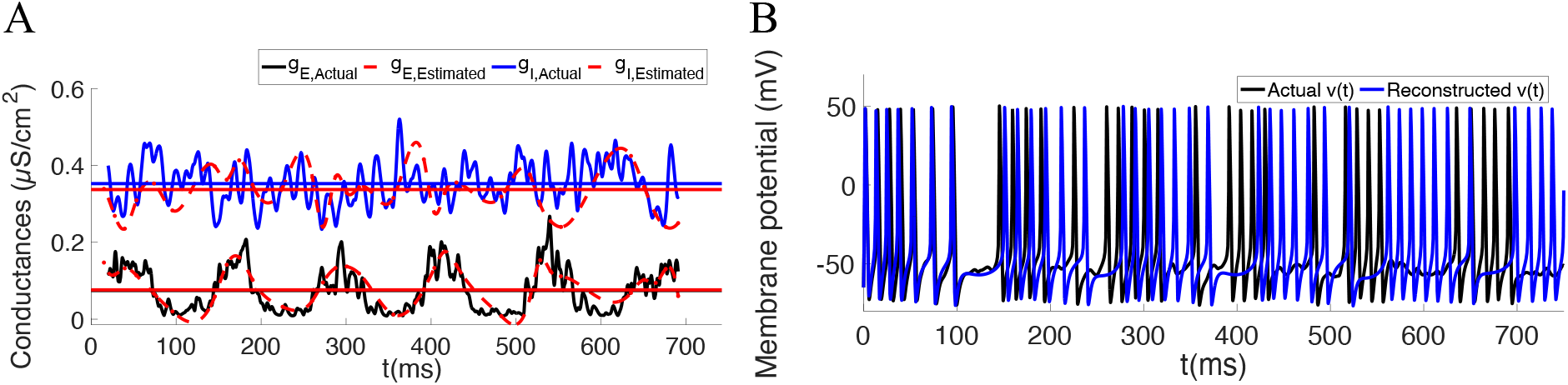
Estimation of an *in sillico* conductances time course from [34]. Panel A shows the actual conductances *g_E_* (black trace) and *g_I_* (blue trace) with the corresponding estimated conductances (dashed red traces). The horizontal lines represent the mean *g_E_* (black line) and *g_I_* (blue line) over time with the corresponding mean also computed from the estimated conductances (red lines). The discretized conductances have been interpolated using a cubic spline interpolation. Panel B shows the actual membrane potential (black trace) and the reconstructed membrane potential (blue trace) which has been computed
considering the estimated conductances in panel A.

## 3 Conclusion and discussion

The estimation procedure presented in this work is, to our knowledge, the first computational method capable of decoupling inhibitory and excitatory synaptic contributions from a single membrane potential trace. It performs robustly across a diverse range of oscillatory conductance profiles, spanning varied frequencies and phase relationships, including in-phase and out-of-phase configurations. Throughout these conditions, the mean squared errors consistently remain below the order of 10^−4^, demonstrating the high efficacy of the estimation method. Although prior knowledge of the base model approximating the isolated target neuron is required, the method adapts to any neural framework as long as its uncoupled activity exhibits periodic oscillations with an input-strength-dependent amplitude. This adaptability provides great flexibility, allowing the procedure to be applied to diverse neuronal types.

Contrary to existing experimental protocols that rely on multiple trials [26], our approach requires neither repetitive measurements nor the heavy infrastructure and operational resources typically needed to acquire such data. Once the base model describing the specific neuron’s activity is established – which may require a single additional recording from the isolated cell – the procedure extracts the time course of both conductances using just one trace. Furthermore, this computational framework exhibits high robustness to measurement noise, effectively minimizing potential errors stemming from experimental instrumentation or trial-to-trial variability. While bypassing complex experimental setups and offering the versatility to adapt to different neural models represent significant advantages, the method remains fundamentally dependent on the accuracy of the underlying base model.

Despite these advantages, when synaptic conductances change abruptly – faster than the timescale of the membrane potential – estimation errors may arise due to the omission of high-frequency components that the algorithm cannot capture. However, the underlying slow oscillations and the time-averaged conductance values remain well predicted. In scenarios where synaptic conductances vary more slowly than the neuronal activity, the estimation accuracy improves further, with mean squared errors dropping to the order of 10^−5^ or even 10^−6^, depending on the phase alignment between the inputs.

Notably, while the excitatory conductance is generally estimated with higher accuracy than the inhibitory one – a phenomenon also reported in [24] for the subthreshold regime – we discovered that this discrepancy is primarily driven by a systematic temporal shift rather than a failure to capture the conductance waveform. By compensating for this displacement through a post-processing cross-correlation analysis, the realignment step drastically minimizes the MSE for *g*_*I*_ near the anti-phase configuration, as well as for both *g*_*E*_ and *g*_*I*_ near the in-phase regime. This adjustment effectively isolates phase delays as a primary source of reconstruction error and brings the accuracy of the inhibitory conductance estimation to a level comparable to that of the excitatory one.

Finally, because these estimation methods are based on point-neuron formulations, they are fundamentally restricted to predicting local synaptic conductances in experimental settings. Under the assumption of linear models, recent advances, such as the ball-and-stick neuron framework proposed by [37], have introduced ways to calculate effective synaptic conductances that mitigate errors arising from space-clamp limitations in neurons with extended dendrites. Combining our non-linear estimation procedure with such multi-compartmental frameworks represents a promising avenue to overcome spatial filtering effects in complex dendritic trees. In addition, while a bijection between (*T, A*) and (*g*_*E*_, *g*_*I*_) seems to hold for our current model (results not shown), alternative base models might lack this uniqueness. To address this potential constraint and prevent computational issues, the algorithm is designed to output a single pair of conductances along with a specific warning if non-bijective regions are encountered.

## Supporting information

## S1 Appendix Mathematical model of the stellate cell.

The stellate cell model is taken from [32]. The membrane potential dynamics is given by

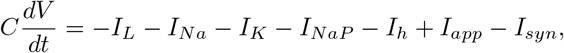

where *C* is the membrane capacitance, *I*_*syn*_ the synaptic current and *I*_*app*_ the applied current. The leakage and ion currents are given by:

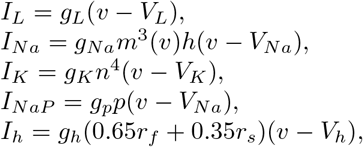

where *V*_*ion*_ and *g*_*ion*_ denote the corresponding ion reversal potential and maximal conductance, respectively. The gating variables *w*, which can be either *m, h, n, p, r*_*f*_ or *r*_*s*_, follow the differential equation

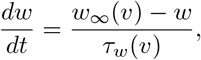

where

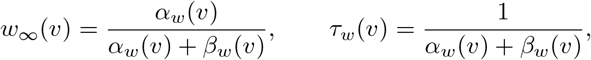

and the gating dynamics are given by:

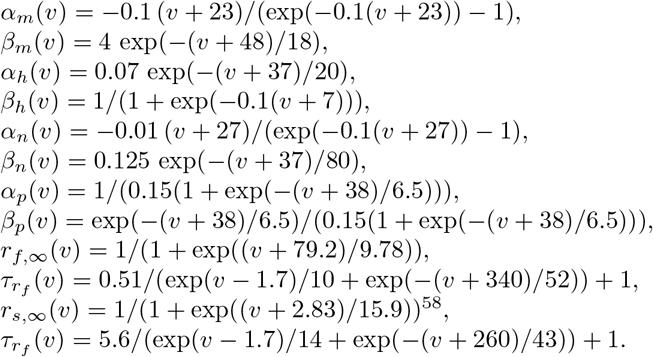

The biophysical parameters are:

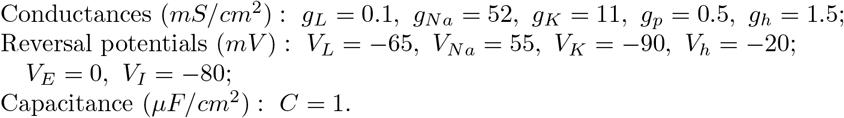

In the absence of synaptic inputs, this model presents spikes for an applied current greater than *I*_*T*_ = − 1.46*µA/cm*^2^. In our case, to have a high frequency of spikes, we inject a constant current equal to *I*_*app*_ = 8 *µA/cm*^2^.

## S2 Appendix Supplementary figures.

## Acknowledgments

AG is supported by grant PID2024-155942NB-I00 funded by MICIU/AEI/10.13039/501100011033 and ERDF, UE, Severo Ochoa and Marí de Maeztu Programs for Centers and Units of Excellence in R& D (CEX2020-001084-M) funded by MCIN/AEI/ 10.13039/501100011033 and by ERDF “A way of making Europe”, and AGAUR grant 2021-SGR-01039. RMDM, AET and CV are support by grant PID2023-151974NB-I00 funded by MICIU/AEI/10.13039/ 501100011033/FEDER, UE. RMDM is also supported by the Conselleria d’Educació i Universitats del Govern de les Illes Balears under grant FPU2025-012-C.

---

**S1 Fig.**
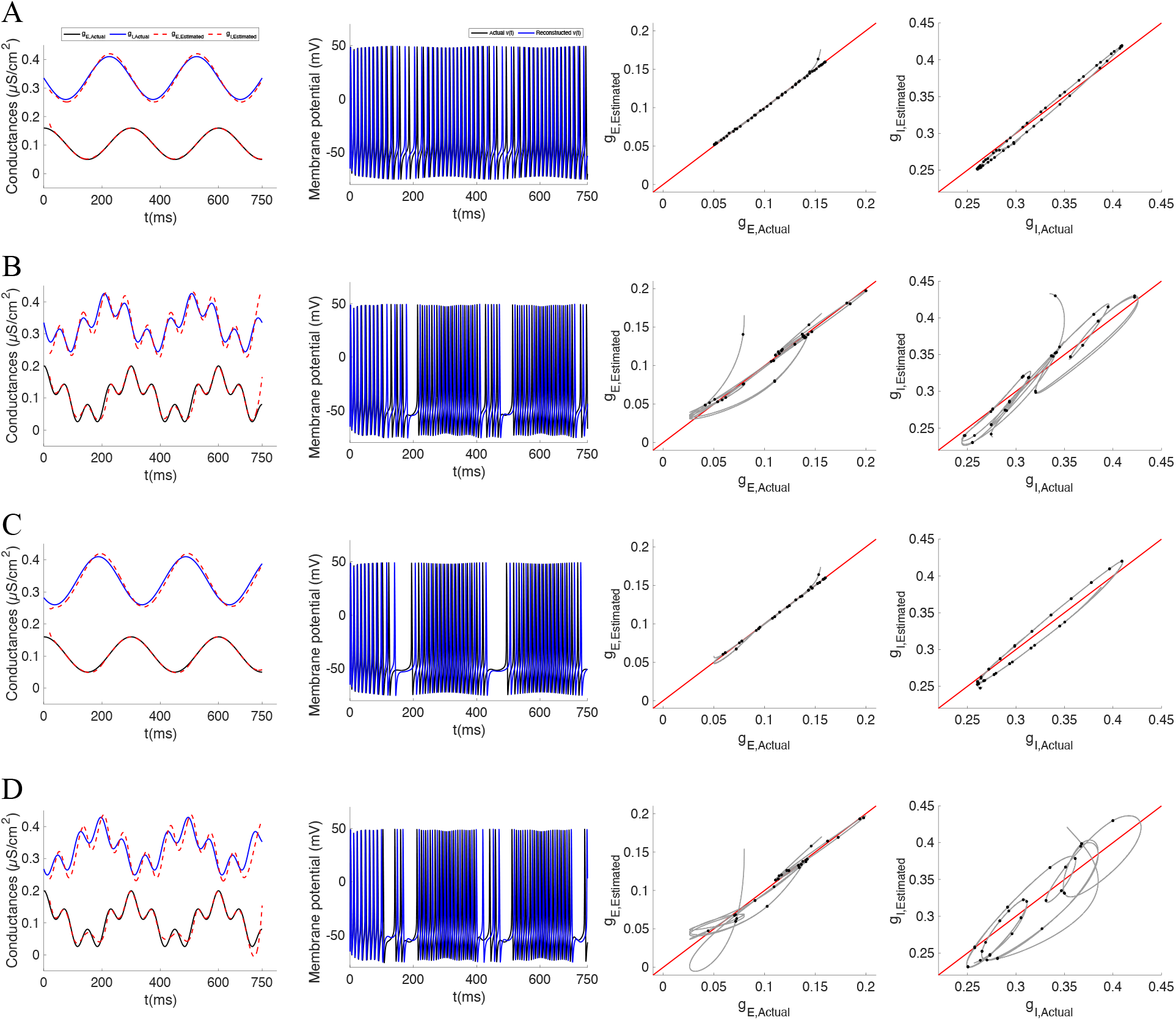
Estimation of the conductances time course when the inhibitory and the excitatory conductances are out-of-phase. Different panels depict the results of the estimation when *g*_*E*_ and *g*_*I*_ have a phase shift of 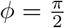 (Panels A and B) or a phase shift of 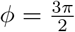 (Panels C and D) considering a single oscillation frequency (Panels A and C) or two (Panels B and D). The equations and rest of parameters for the conductances in rows A and C are provided in equation (4) while those for rows B and D are provided in equation (5). From left to right, the different columns depict for: (1) the actual conductance traces (excitatory conductance in black and the inhibitory one in blue) and the corresponding estimated conductances provided by the estimation procedure (dashed red lines); (2) the actual membrane potential trace (black trace) and the reconstructed one by considering the estimated conductances (blue trace); (3) the actual excitatory conductances trace versus the estimated ones; and (4) the actual inhibitory conductances versus the estimated ones. In the 3th and 4th columns, black dots are the considered values in the estimation, while the grey trace represents the interpolation of these values (the traces shown in the panel of corresponding 1st column). In all panels of the 3rd and 4th columns, the red line represents the identity line, the dots correspond to the discretized estimated conductances while the black traces show the conductances after interpolate them using the cubic spline interpolation.

**S2 Fig.**
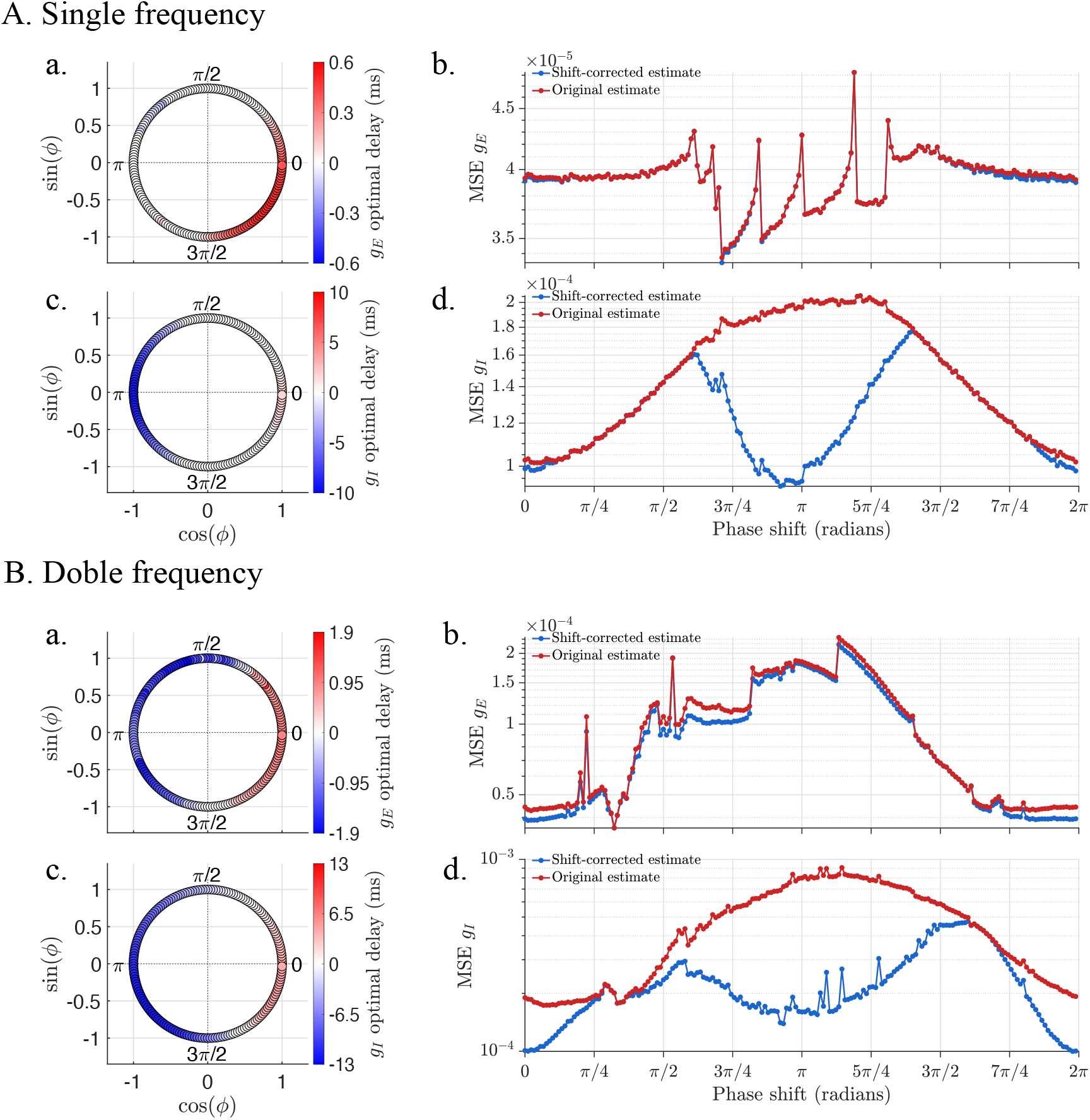
Effect of phase shift between *g*_*E*_(*t*) and *g*_*I*_ (*t*) on oscillatory synaptic conductance estimation under the presence of noise (*σ* = 0.01). Panels A and B show the results when the excitatory and the inhibitory conductances are composed of a single or two oscillatory frequency components, respectively. At each panel, first column depicts the optimal temporal offsets, *τ*_opt,*E*_ (subpanels A.a and B.a) and *τ*_opt,*I*_ (subpanels A.c and B.c, that maximizes the correlation between the estimated and original excitatory (*g*_*E*_) and inhibitory (*g*_*I*_) synaptic conductances as a function of the imposed phase shift. Here, positive (red) and negative (blue) values indicate that the estimated conductance must be advanced or delayed, respectively, to achieve the best alignment. The second column represents the mean squared error (MSE) of the *g*_*E*_ (subpanels A.b and B.b) and *g*_*I*_ (subpanels A.d and B.d) reconstructions, before (original estimate, in red) and after (shift-corrected estimate, in blue) temporal alignment using the optimal offset shown in the first column. In all panels, the phase shifts *ϕ* are evaluated across the full range [0, 2*π*).

**S3 Fig.**
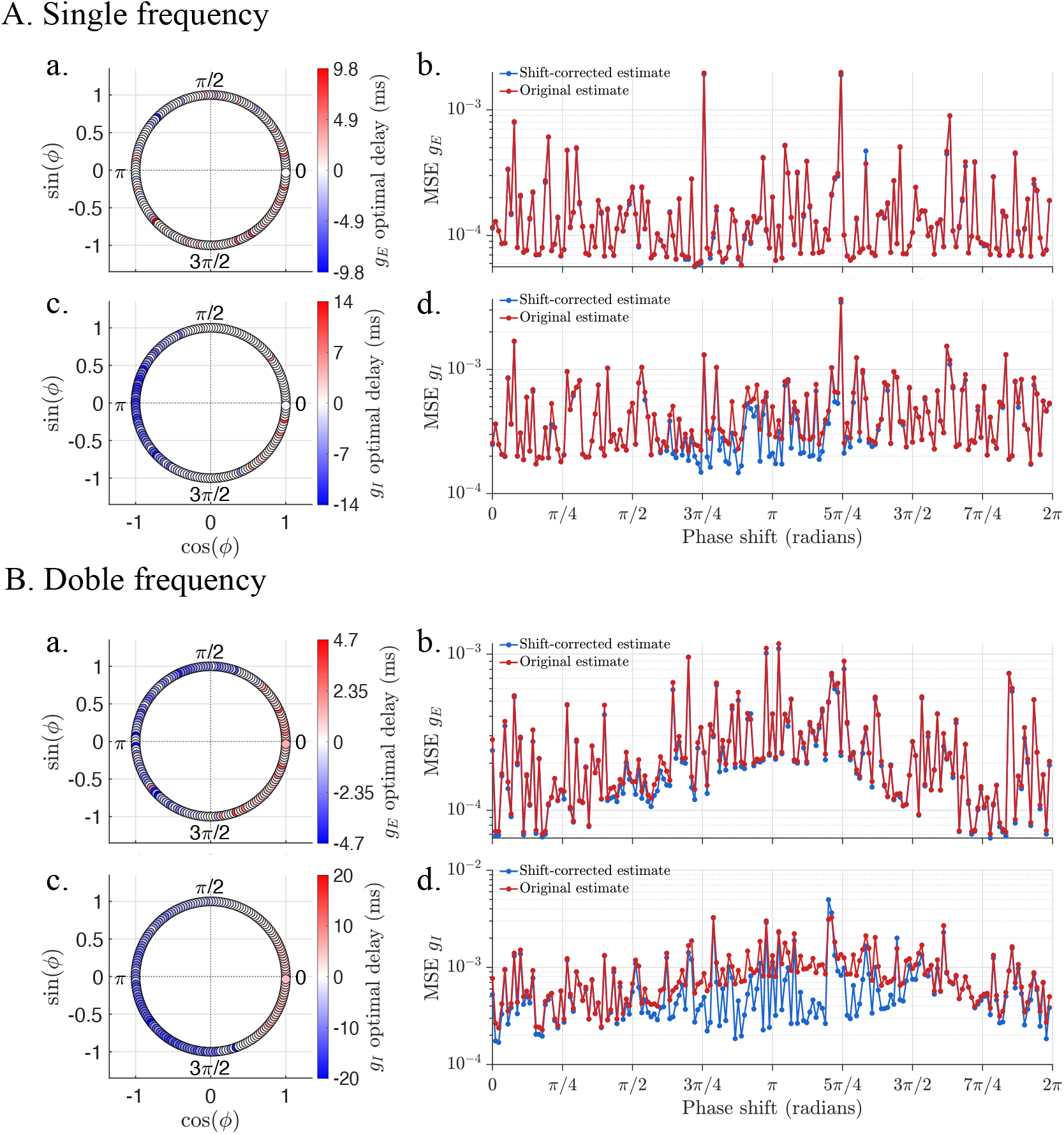
Effect of phase shift between *g*_*E*_(*t*) and *g*_*I*_ (*t*) on oscillatory synaptic conductance estimation under the presence of noise (*σ* = 0.1). Panels A and B show the results when the excitatory and the inhibitory conductances are composed of a single or two oscillatory frequency components, respectively. At each panel, first column depicts the optimal temporal offsets, *τ*_opt,*E*_ (subpanels A.a and B.a) and *τ*_opt,*I*_ (subpanels A.c and B.c, that maximizes the correlation between the estimated and original excitatory (*g*_*E*_) and inhibitory (*g*_*I*_) synaptic conductances as a function of the imposed phase shift. Here, positive (red) and negative (blue) values indicate that the estimated conductance must be advanced or delayed, respectively, to achieve the best alignment. The second column represents the mean squared error (MSE) of the *g*_*E*_ (subpanels A.b and B.b) and *g*_*I*_ (subpanels A.d and B.d) reconstructions, before (original estimate, in red) and after (shift-corrected estimate, in blue) temporal alignment using the optimal offset shown in the first column. In all panels, the phase shifts *ϕ* are evaluated across the full range [0, 2*π*).

